# IL-33 protects from recurrent *C. difficile* infection by restoration of humoral immunity

**DOI:** 10.1101/2024.11.16.623943

**Authors:** Farha Naz, Nicholas Hagspiel, Mary K. Young, Jashim Uddin, David Tyus, Rachel Boone, Audrey C. Brown, Girija Ramakrishnan, Isaura Rigo, Gregory R. Madden, William A. Petri

## Abstract

Clostridioides difficile infection (CDI) recurs in one of five patients. Monoclonal antibodies targeting the virulence factor TcdB reduce disease recurrence, suggesting that an inadequate anti-TcdB response to CDI leads to recurrence. In patients with CDI, we discovered that IL-33 measured at diagnosis predicts future recurrence, leading us to test the role of IL-33 signaling in the induction of humoral immunity during CDI. Using a mouse recurrence model, IL-33 was demonstrated to be integral for anti-TcdB antibody production. IL-33 acted via ST2+ ILC2 cells, facilitating germinal center T follicular helper (GC-Tfh) cell generation of antibodies. IL-33 protection from reinfection was antibody-dependent, as μMT KO mice and mice treated with anti-CD20 mAb were not protected. These findings demonstrate the critical role of IL-33 in generating humoral immunity to prevent recurrent CDI.

Graphical Abstract:
IL-33 restoration induces toxin-B-specific antibody production via the ILC2-TFH axis. In the left panel, IL-33 remediation increases ILC2s, subsequently inducing TFH directly or indirectly. TFH cell induction is pivotal for the production of antibodies. IL-33 also downregulates type 1 and type 3 immunity, favoring type 2 immunity to enhance host survival and reduce morbidity. The middle panel illustrates antibiotic-induced dysbiosis, resulting in decreased IL-33 levels and reduced antibody production. The right panel demonstrates the protective effect in reinfection, attributed to toxin-specific antibodie generated by IL-33 remediation.

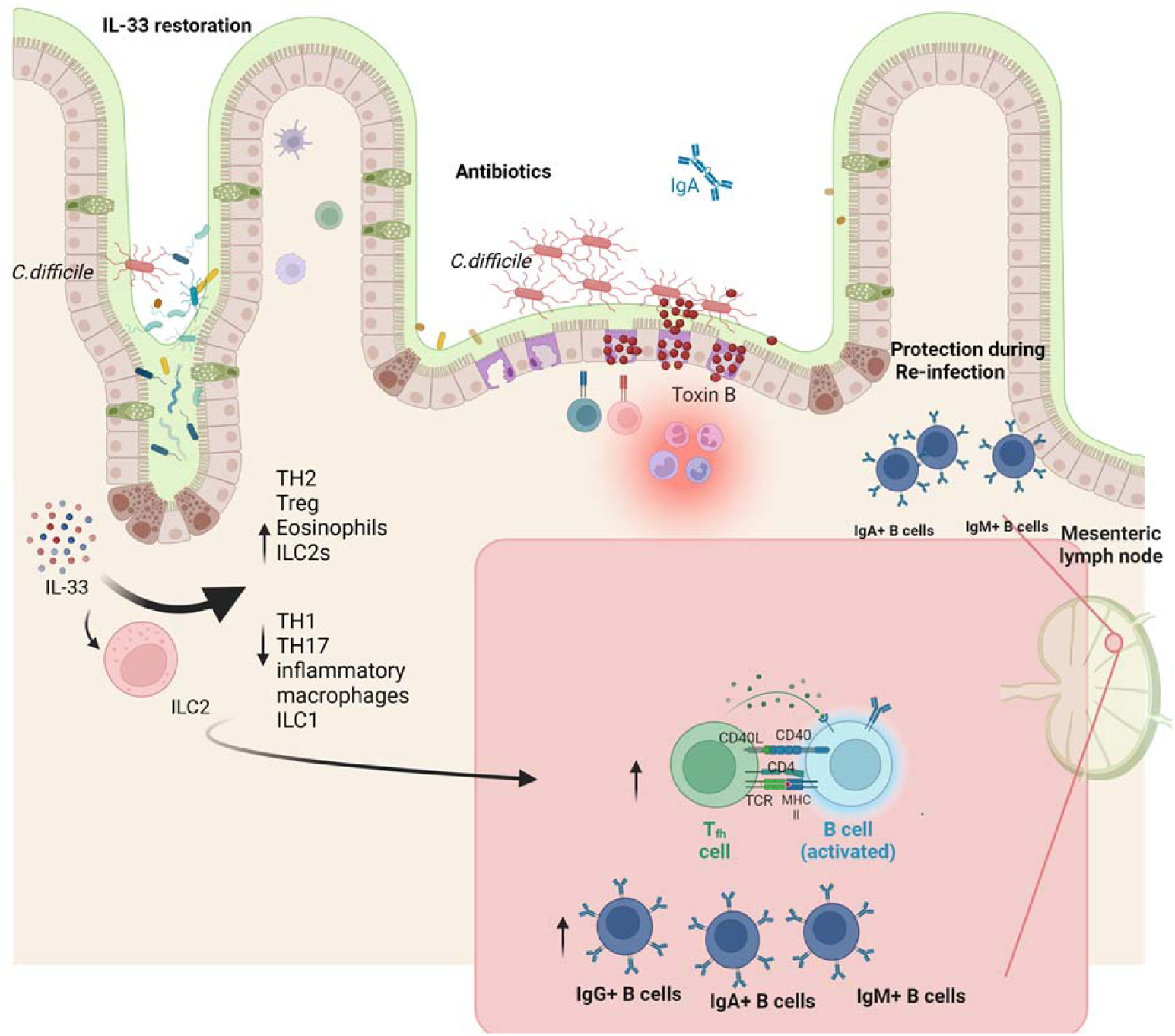

## Introduction

A unique and challenging aspect of *Clostridioides difficile* infection (CDI) is its tendency to recur in up to 25% of patients, a risk that increases with each subsequent recurrent episode(1, 2). Antibiotics targeting *C. difficile* are a double-edged sword in CDI, which despite treating the acute infection further disrupts the intestinal microbiome, predisposing patients to recurrence(3). The most effective therapy to prevent recurrent CDI (rCDI) is fecal microbiota transplantation (FMT) and newer microbiota therapeutics (e.g., SER109, RBX2660); however, these are not 100% efficacious and have significant limitations (i.e., cost/logistical barriers and the risk of transmitting pathogens in the case of FMT)(4, 5).

Recurrent infection can partly be attributed to compromised adaptive immunity, suggesting a role for the immune system in preventing and managing repeated infections(6). *C. difficile* toxin B plays a key role in CDI pathogenesis(7-9), and exogenous IgG antibodies against toxin B are capable of averting rCDI(10, 11). Investigation into the role of immunoglobulins IgA, IgG, and IgM has been a consistent focus in studies involving human CDI patients(10). For example, reduced levels of antibodies (IgG, IgA) against TcdA and TcdB in the serum were linked to recurrence, whereas antibodies targeting cell surface antigens did not show any such correlation(12). Recent findings indicate that the presence of toxin B-specific IgG during acute CDI correlates with a delay in the onset of recurrence (11, 13).

Previous work has shown that IL-33 prevents mortality and epithelial disruption by activating ILC2s in the acute mouse model of CDI(14). The microbiota influences IL-33 expression, and dysregulated IL-33 signaling predicts acute *C. difficile*-associated mortality in humans(14), emphasizing its crucial role in the defense against acute CDI. In this investigation, it is demonstrated that the type 2 alarmin IL-33 serves as a biomarker of recurrence in humans. Additionally, utilizing a male mouse model, the pivotal role of IL-33 in antibody-dependent protection from recurrence is elucidated.

## Results

### IL-33-induced toxin-specific antibodies in the *C. difficile* mouse model

Studies have shown that IL-33 triggers the activation of ILC2s(14), which has the potential to enhance humoral immunity(15). Antibodies to *C. difficile* toxin B are known to prevent recurrence(10). We hypothesized that IL-33 promotes ILC2-dependent anti-toxin antibody production. In the mouse model of acute (primary) CDI, antibiotics induce susceptibility by decreasing IL-33 and subsequent IL-33 activation of ILC2^16^. The antibiotic-induced deficiency in IL-33 in acute CDI could therefore predispose to recurrent infection by impairing the production of anti-toxin B antibody.

To investigate the role of IL-33 in anti-toxin B antibody production, IL-33 was first supplemented in the acute CDI mouse model. Prior to infection with the hypervirulent epidemic R20291 strain, mice were given antibiotics, and IL-33 protein was administered daily for five days by intraperitoneal injection (0.75 μg/mouse) **(Fig. 1A).** IL-33 treatment reconstituted the antibiotic-depleted IL-33 protein level within the colon before infection (**Supplementary Fig. 1A).** As previously observed, acute CDI was less severe in IL-33-treated mice as shown in the survival curve, weight loss, and clinical scores between the groups (**Figs. 1B, 1C and 1D**) (14). Toxin B-specific antibodies (IgG, IgM, IgA) at 15 days post-infection were higher in cecal contents and plasma of IL-33-treated mice (**Figs. 1E, 1G, and 1H**). A similar IL-33 induction of anti-toxin B antibody (IgG) was seen after infection with the classical *C. difficile* strain VPI 10463 (**Fig. 1F**). Survival, weight loss, and clinical scores for VPI strain infection were found similar to those of R20291 (data not shown). *C. difficile* burden at day 15 post-infection was unaltered by IL-33 (**Supplementary Figs. 1B and 1C).** It is concluded that the administration of IL-33 at the time of antibiotic pretreatment protects from acute CDI and enhances the production of anti-toxin antibodies.

**Fig 1:**
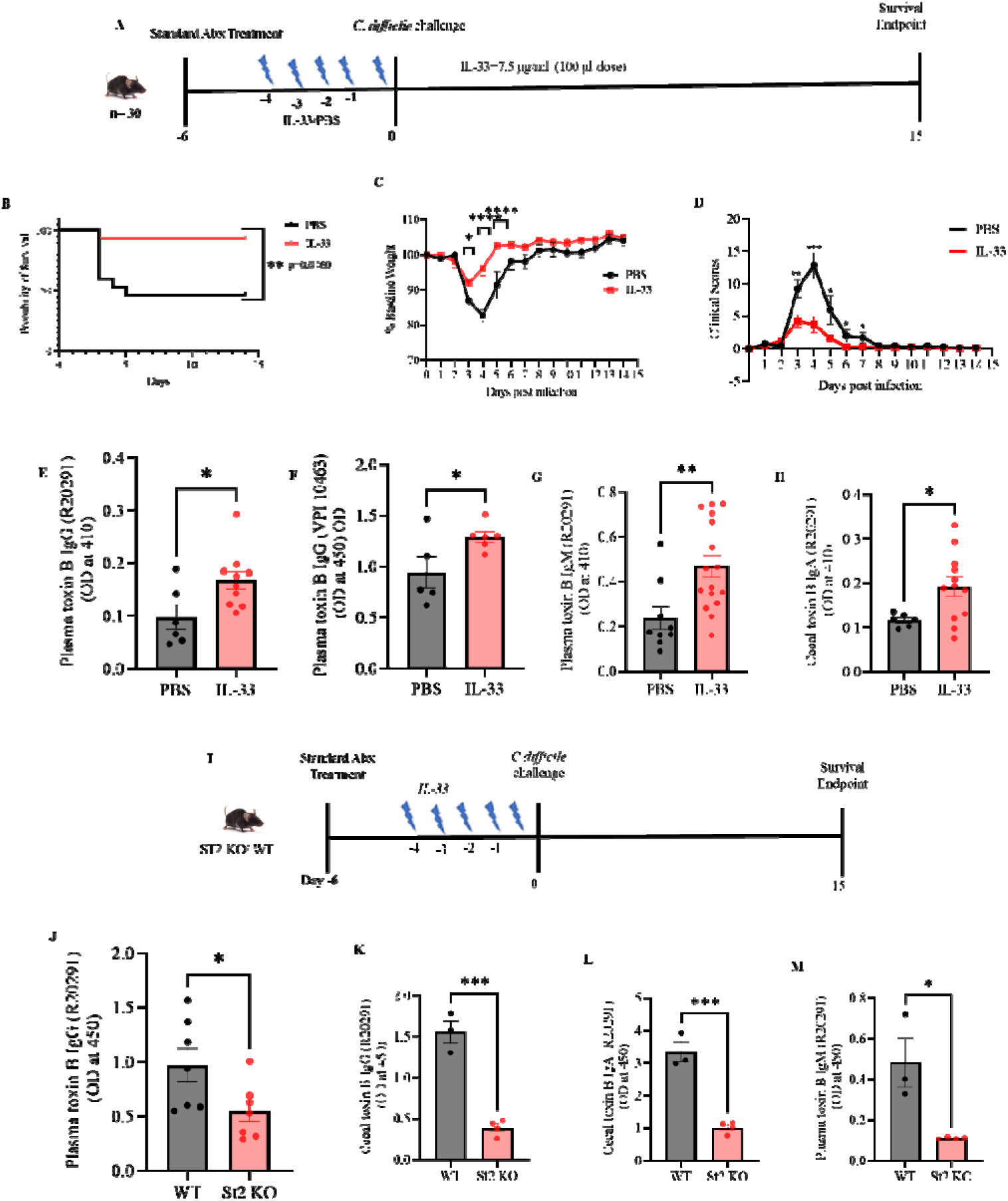
IL-33 increases toxin-specific antibody in mice after first infection with *C. difficile*. IL-33 (0.75 µg) was administered i.p. on days -4 to 0 to C57BL/6J mice (A–H) and/or ST2−/− mice (I-M). Mice were infected with *C. difficile* strain R20291 (A-E and G-M) or VPI 10463 (F). On post-infection day 15 antibodies were measured in plasma and cecal content. (A) Schematic diagram showing infection and treatment timeline; (B) survival curves; (C) weight loss; (D) clinical scores; (E) plasma toxin B specific IgG from mice infected with *C. difficile* strain R20291; (F) plasma toxin B specific IgG from mice infected with *C. difficile* strain VPI 10463; (G) plasma toxin B specific IgM from mice infected with *C. difficile* strain R20291; (H) cecal content toxin B specific IgA from mice infected with *C. difficile* strain R20291. (I-M) WT vs ST2-/- mice infected with *C. difficile* strain R20291: (I) Schematic diagram showing infection and treatment timeline; (J) plasma IgG; (K) cecal IgG; (L) cecal IgA; and (M) plasma IgM. B, Comparison made by log-rank test ( n = 30 in both groups). C, D, Comparison made by two-tailed Student’s t-test (C, D n = 30). E, F, G, H, J, K, L, M, A two-tailed t-test for normally distributed data and a Mann-Whitney test for non-normally distributed data were used. *P < 0.05, **P < 0.01, and ***P < 0.001. The error bar indicates SEM.

The ST2 receptor for IL-33 is expressed on many immune cells including B cells(16). To test if the induction in antibody production was due to IL-33 signaling via its receptor ST2, the ST2 knockout, and wild-type mice were utilized to compare antibody production after IL-33 administration **(Fig. 1I).** Cecal contents and plasma were collected 15 days post-infection. As expected, IL-33 did not induce anti-TcdB antibodies (IgG, IgA, IgM,) production in ST2 KO mice **(Figs. 1J, 1K, 1L, and 1M).** This led to the conclusion that IL-33 exerts its effects through the ST2 receptor.

### Decrease in severity of *C. difficile* reinfection by IL-33

To test if restoration of IL-33 during acute CDI could protect from reinfection, a murine model of reinfection was utilized (**Fig. 2**). C57BL/6J mice were infected on day 0 with *C. difficile* strain R20291 after having been pretreated with antibiotics with or without administration of exogenous IL-33. First, we established the mouse reinfection model of *C. difficile*. Of note, we did not find any difference in bacterial colonization between the groups throughout the infection trajectory (data not shown). After recovery from the primary infection, on day 54, the mice retreated with antibiotics before reinfection with 10^4^ *C. difficile* spores from strain R20291 (**Fig. 2A**). Antibiotic retreatment was important to clear the bacterial colonization as even after the 100 days of primary infection, *C. difficile* colonization and toxin production were observed by using kits (Toxin B in the stool was detected using the ELISA kit (TechLab Inc., catalog #T5015). The number of bacteria was measured by Tech lab kit (TechLab Inc., catalog #TL5025). Unlike the control mice, mice treated with IL-33 during the primary acute CDI did not show clinical signs or lose weight upon reinfection (**Figs. 2B and 2C**). Further, IL-33 treatment during acute CDI led to improved gut barrier function during reinfection (**Fig. 2D**). The treated group also experienced reduced submucosal edema and epithelial damage (**Figs. 2E and 2F**). It is concluded that IL-33 restoration during primary CDI promoted gut integrity to protect from recurrent infection.

**Fig 2:**
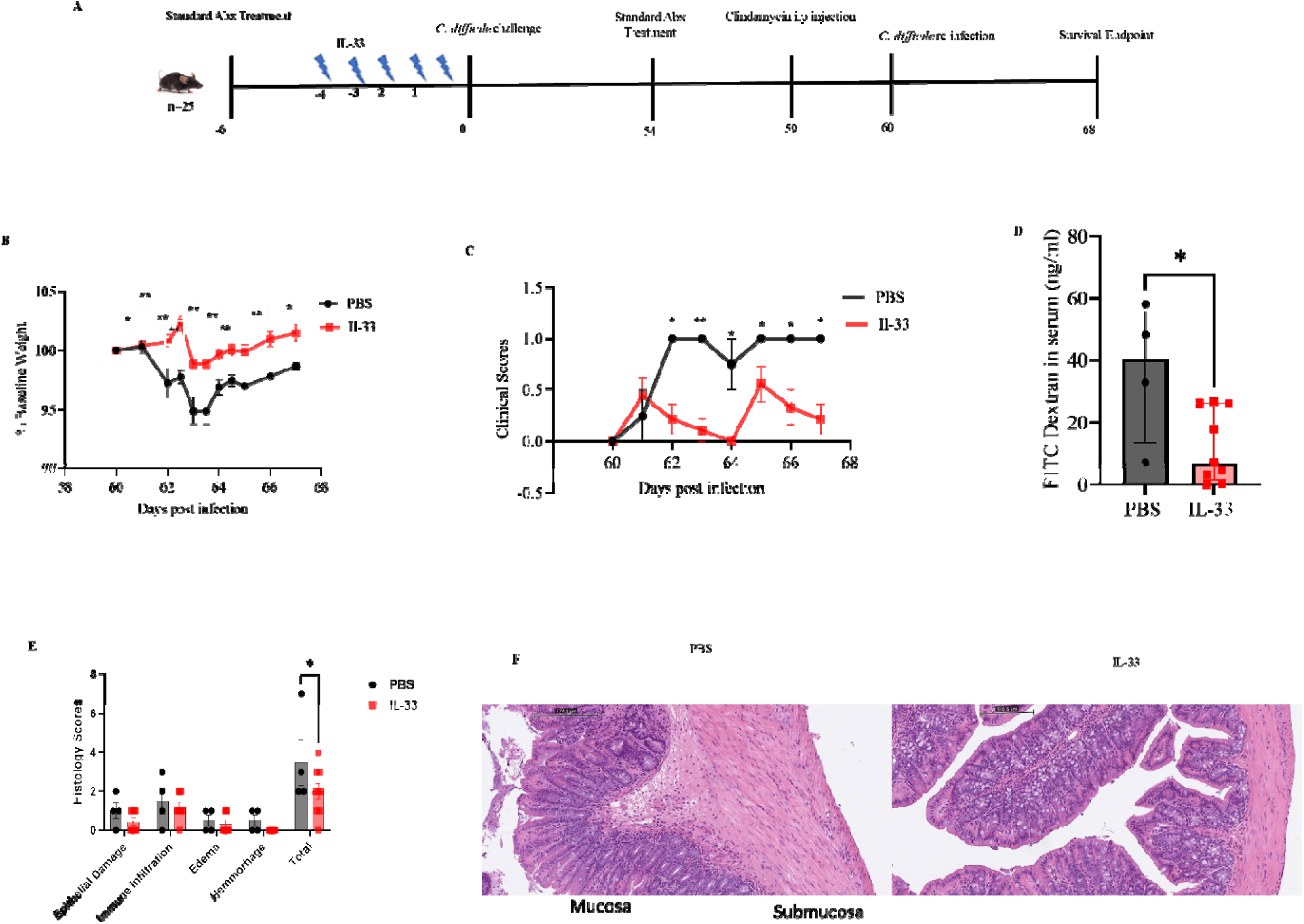
IL33 protects from a 2nd C. difficile infection. IL-33 (0.75 µg) was administered i.p. on days -4 to 0 and wild-type mice were infected on day 0 and again on day 60 with C. difficile strain R20291. (A) Experimental design for 2nd infection; (B) 2nd infection weight loss; (C) clinical scores; (D) FITC-dextran gut permeability test; (E) epithelial damag scoring; (F) representative H&E stain of the colon. C, D, Comparison made by two-tailed Student’s t-test (C, D n = 13). D, Mann-Whitney test for non-normally distributed data was used E, Šídák’s multipl comparisons test was used to determine the statistical significance between groups. Statistical significance is demarked as *P < 0.05, **P < 0.01, and ***P < 0.001. The error bar indicates SEM in B, D, and E but D indicates the median with interquartile range.

We then tested if IL-33 could be used after a primary infection to prevent recurrence. IL-33 was administered after the acute infection prior to rechallenge **(Supplemental Fig. 2A)**. The group who received IL-33 before reinfection regained weight faster than the PBS recipient group **(Supplemental Fig. 2B)** and returned to a clinical score of zero faster than the PBS group **(Supplemental Fig 2C)**. The IL-33-treated group also produced more IgM and IgG in the serum than the control group **(Supplemental Figs. 2D and 2E)** suggesting that protection from reinfection may be achieved through increased toxin-specific antibody production. Of note, IL33 did not alter the colonization of the bacteria.

### Importance of antibody production for IL-33 mediated protection from reinfection

In order to determine if IL-33-mediated protection against reinfection was mediated by antibody, μMT knockout mice that lack mature B-cells were pretreated with IL-33 and infected with *C. difficile* (**Fig. 3A** During the first ten days of the initial *C. difficile* infection, there was no difference in weight loss or clinical scores between wild-type (WT) and μMT knockout mice, consistent with prior work that showed no role of B cells and T cells in the acute phase of CDI (**Figs. 3B and 3C)**(14, 17). Interestingly, from day 11 onwards, WT mice gained significantly more weight than the μMT knockout mice, suggesting a role of antibodies in the sub-acute recovery phase of primary CDI. To confirm the absence of antibodies, plasma and stool IgG, IgM, and IgA were checked from the μMT knockout mice that did not produce toxin B-specific antibodies (**Supplementary Figs. 3A, 3B, 3C, 3D, 3E**) (18). Mice were retreated with antibiotics cocktails and reinfected on day 60 after the primary *C. difficile* infection. The μMT knockout mice lost more weight than the wild-type mice when given antibiotics and reinfection (**Fig. 3D**). Interestingly, μMT knockout mice had higher levels of toxin A/B in the stool (**Fig. 3E**), had increased gut permeability calculated by FITC-dextran gut permeability assay (**Fig. 3F**), and greater submucosal edema and epithelial damage (**Figs. 3G and 3H**). Sample were collected after the endpoint of the experiment i.e. post 11 days after reinfection. Surprisingly, μMT knockout mice had a lower *C. difficile* bacterial burden as measured by a GDH ELISA kit (TechLab Inc., catalog #TL502) (**Supplementary Fig. 3F**).

**Fig 3:**
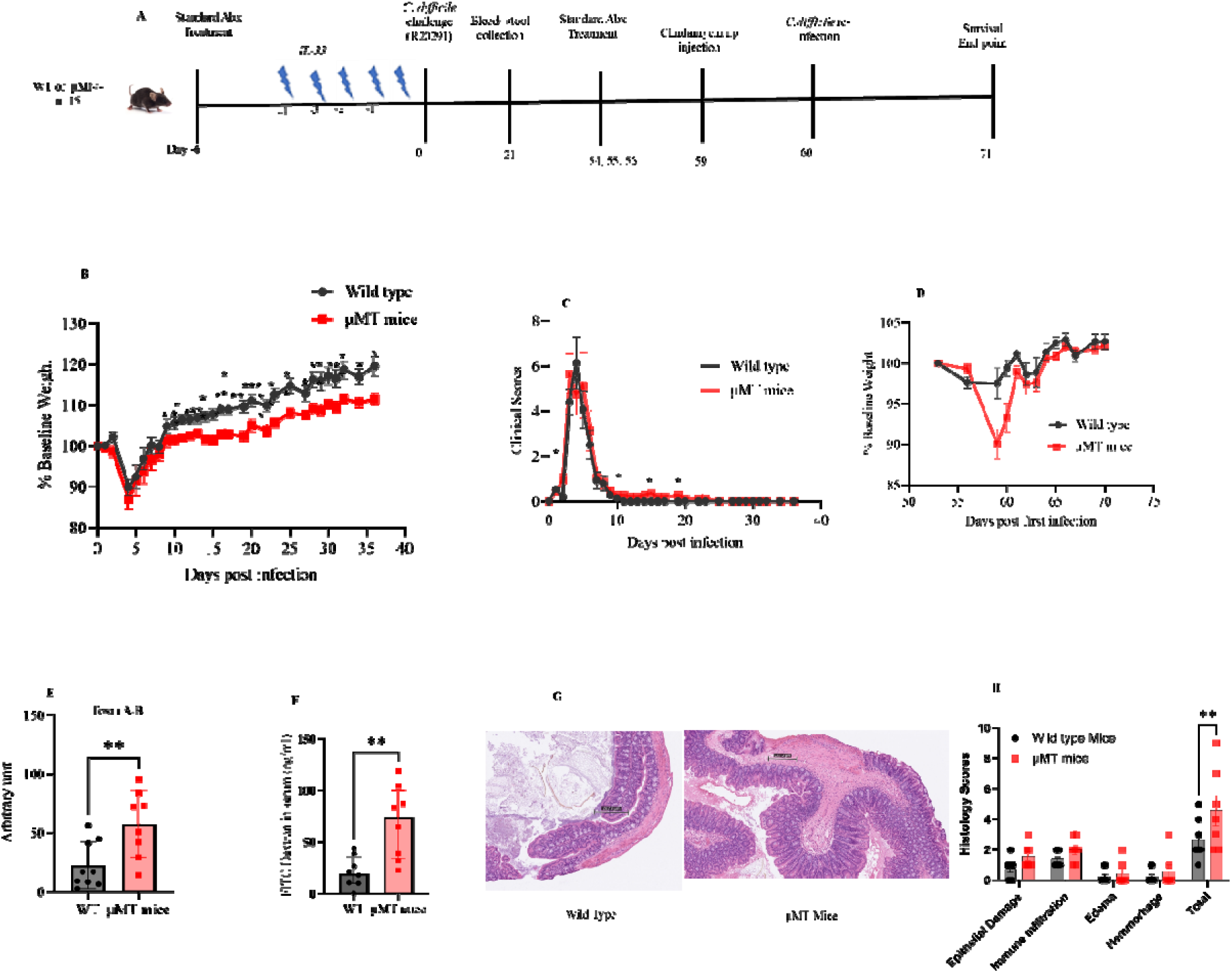
IL33 protection from a 2nd C. difficile infection is antibody-dependent. WT and μMT KO mice were administered IL-33 (0.75 µg) i.p. on days -4 to 0 and mice infected on day 0 and reinfected on day 60 with C. difficile strain R20291. (A) Experimental design; (B) 1st infection weight loss; (C) clinical scores. WT and μMT KO mice were reinfected with C. difficile R20291 60 day after the first infection. Reinfection (D) weight loss; (E) stool C. difficile toxin A and B; (F) FITC-dextran gut permeability assay; (G) H&E stain; (I) epithelial damage scoring. B, C, D, Comparison made by two-tailed Student’s t-test (B, C n = 30, D n=18). E, a two-tailed t-test was used and the error bar indicates SEM. F, a Mann-Whitney test was used and the error bar indicates the median with interquartil range. H, Šídák’s multiple comparisons test was used. Statistical significance is demarked as *P < 0.05, **P < 0.01, and ***P < 0.001.

Given the limitations of μMT knockout mice in producing IgE and IgG antibodies(19), we employed αCD20 to deplete B cells (**Fig. 4A**). During acute CDI, the αCD20 treated mice had slightly higher mortality (**Fig. 4B**) but no difference in weight loss or clinical scores was found **(Figs. 4C and 4D).** Upon reinfection, the αCD20 treated mice lost more weight (**Fig. 4E**), had higher clinical scores (**Fig. 4F**), and worse submucosal edema and epithelial damage (**Figs. 4G and 4H**). No toxin-specific antibodies (IgG, IgA) were detected at the end of reinfection estimated from plasma and cecal tissue **(Figs. 4I, 4J, 4K, and 4L**). B cell depletion was confirmed in the colon and MLN **(Figs. 4M and 4N & 4V and 4W**). Interestingly, we observed that TH2 **(Figs. 4O& 4P)** and Treg cell populations **(Figs. 4S & 4U)** in the colon were significantly lower, and neutrophils were higher **(Figs. 4Q and 4R**) in the αCD20 treated mice. That itself described more inflammation in the colon of αCD20 treated mice. The overall conclusion was drawn that antibody production was required for IL-33-mediated protection from recurrent *C. difficile*.

**Fig 4:**
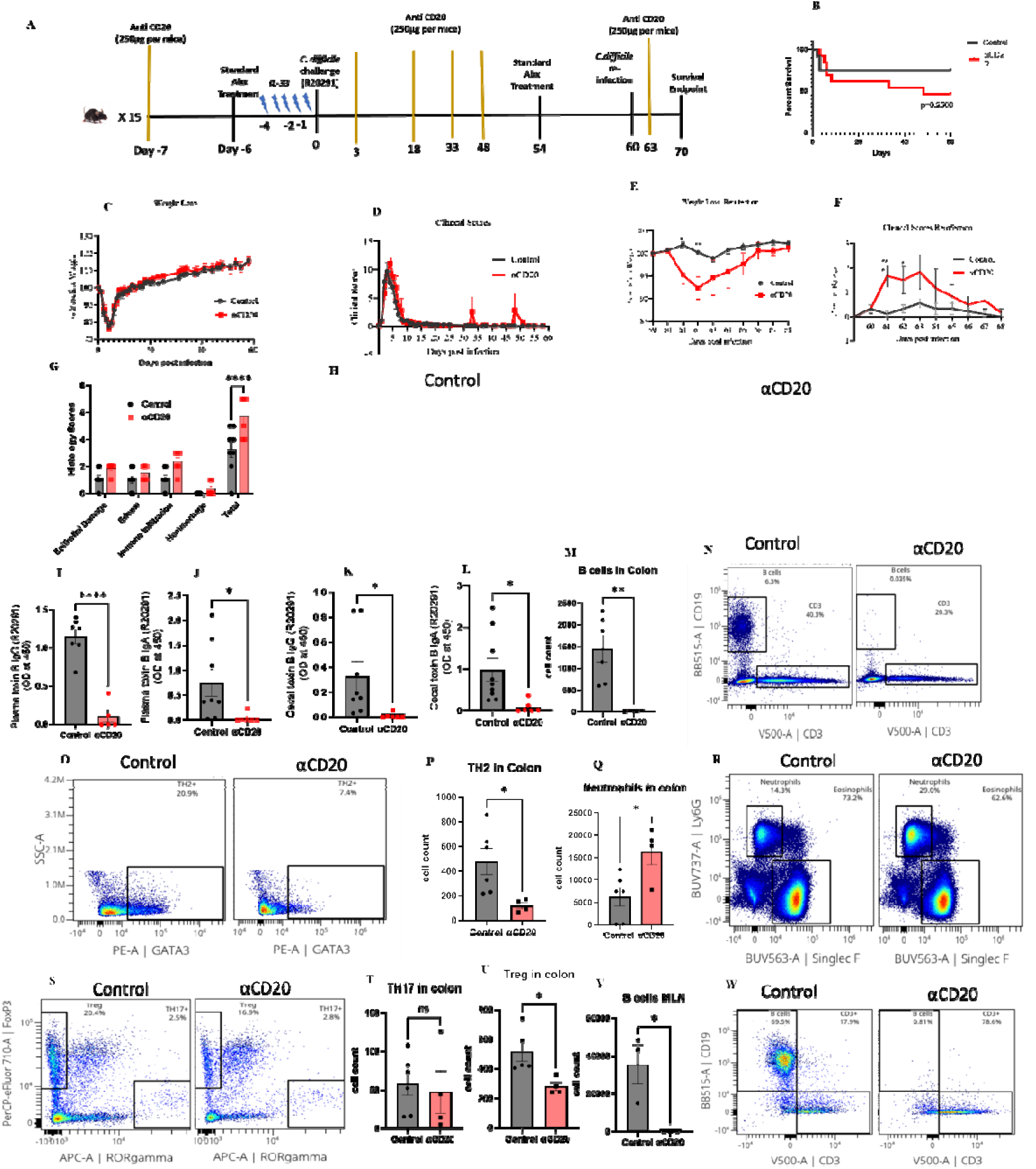
Antibody deficient mice (**α**CD20 treated) lost IL-33 mediated protection from a 2nd C. difficile infection. αCD20 was administered to deplete B cells on days -7, 3, 18, 33, 48, and 63, and mice infected on day 0 and 2nd time infected on day 60 with C. difficile strain R20291. (A) Experimental design; (B) survival curve; (C) 1st infection weight loss; (D) 1st infection clinical scores; (E) reinfection weight loss; (F) reinfection clinical scores (G) epithelial damage scoring; (H) H&E stain. (I) toxin B specific plasma IgG; (J) IgA; (K) cecal IgG; and (L) cecal IgA measured on day 10 post 2nd infection. MLN and colon were harvested on day 10 post-reinfection. Colonic (M, N) B cells ( CD45+ CD3-CD19+); (O, P) TH2 cells (CD45+CD3+ CD4+ GATA3+); (Q, R) neutrophils; (S, T, U) Treg (CD45+CD3+ CD4+ FOXP3+)and TH17(CD45+CD3+ CD4+ RORγt+) cells; and (V, W) MLN B cells. B, Comparison made by log-rank test ( n = 26 in both groups). C, D, E, F Comparison made by two-tailed Student’s t-test (C, D n = 26, E, F n=15). G, Šídák’s multiple comparisons test was used. I, J, K, L, M, P, Q, T, U, V A two-tailed t-test for normally distributed data and a Mann-Whitney test for non-normally distributed data were used. Statistical significance is demarked as *P < 0.05, **P < 0.01, ***P < 0.001, and ****P < <0.0001. The error bar indicates SEM.

The next question was whether IL-33-mediated protection from recurrent *C. difficile* was strain-specific. The hypervirulent strain R20291 produces a toxin B (TcdB2) antigenically distinct from the TcdB1 of the classical strain VPI 10463. The expectation was that if IL-33 protection from recurrent R20291 infection was due to anti-TcdB2 antibody production, IL-33 would not prevent recurrence from the classical VPI 10463 that produces TcdB1. After a primary infection with *C. difficile* strain R20291 with IL-33, on day 40, mice were reinfected with either R20291 or VPI 10463 (**Supplemental Fig. 4A**). Mice that were reinfected with a different strain (i.e. VPI 10463) than the strain used in a primary infection (i.e. R20291) lost more weight (**Supplemental Fig. 4B**) and had a more severe clinical score (**Supplemental Fig. 4C)** than the group that was reinfected with the same strain, i.e. R20291). It is concluded that the strain specificity of IL-33 mediated protection from recurrence was consistent with its mediation by strain-specific anti-toxin B antibodies.

### IL-33-mediated increase in mucosal Type 2 immunity during primary *C. difficile* infection

To further understand cellular dynamics and the overall host response to the IL-33 treatment in dysbiotic mice before and after the first *C. difficile* challenge, immunophenotyping was done on the host’s innate and adaptive immune response. First, the immune population in infected and noninfected groups was checked after the recovery phase (day 16 post-infection). Neutrophils and TH17 cells were significantly higher in the *C. difficile-infected* mice even after 16 days of infection indicating type 3 immunity dominates and persists beyond the resolution of CDI **(Supplemental Fig. 5)**. During primary CDI, upon IL-33 treatment, ILC2s were increased, and ILC1 and ILC3 populations decreased in the mesenteric lymph nodes (MLN) (**Figs. 5A, 5B, 5C, and 5D**) and colon **(Supplemental Figs. 6A, 6B, 6C and 6D)**. Additionally, there was an increase in TH2 and a decrease in TH1 cell populations before and after the *C. difficile* challenge in MLN (**Figs. 5E, 5F, 5G, 5H, 5I, 5J, 5K, 5L, 5M, 5N, 5O, and 5P**) and colon (**Supplemental Figs. 6E, 6F)** upon IL-33 remediation of dysbiosis. Flow cytometry at days 2 and 6 after primary CDI demonstrated an IL-33-induced downregulation of TH17 cells and upregulation of Treg cells in MLN (**Figs. 6A, 6B, 6C, 6D, 6E, 6F, 6G, 6H, 6I**) and colon (**Supplemental Figs. 6G, 6H, and 6I**). In line with our previous study, IL-33 increased colonic eosinophils and decreased inflammatory monocytes (Ly6C high populations) (**Supplemental Figs. 6J, 6K, 6L, and 6M**) (14). We concluded that IL-33 treatment during acute CDI promoted a long-lasting innate and adaptive type 2 immune response in the intestine and MLN.

**Fig 5:**
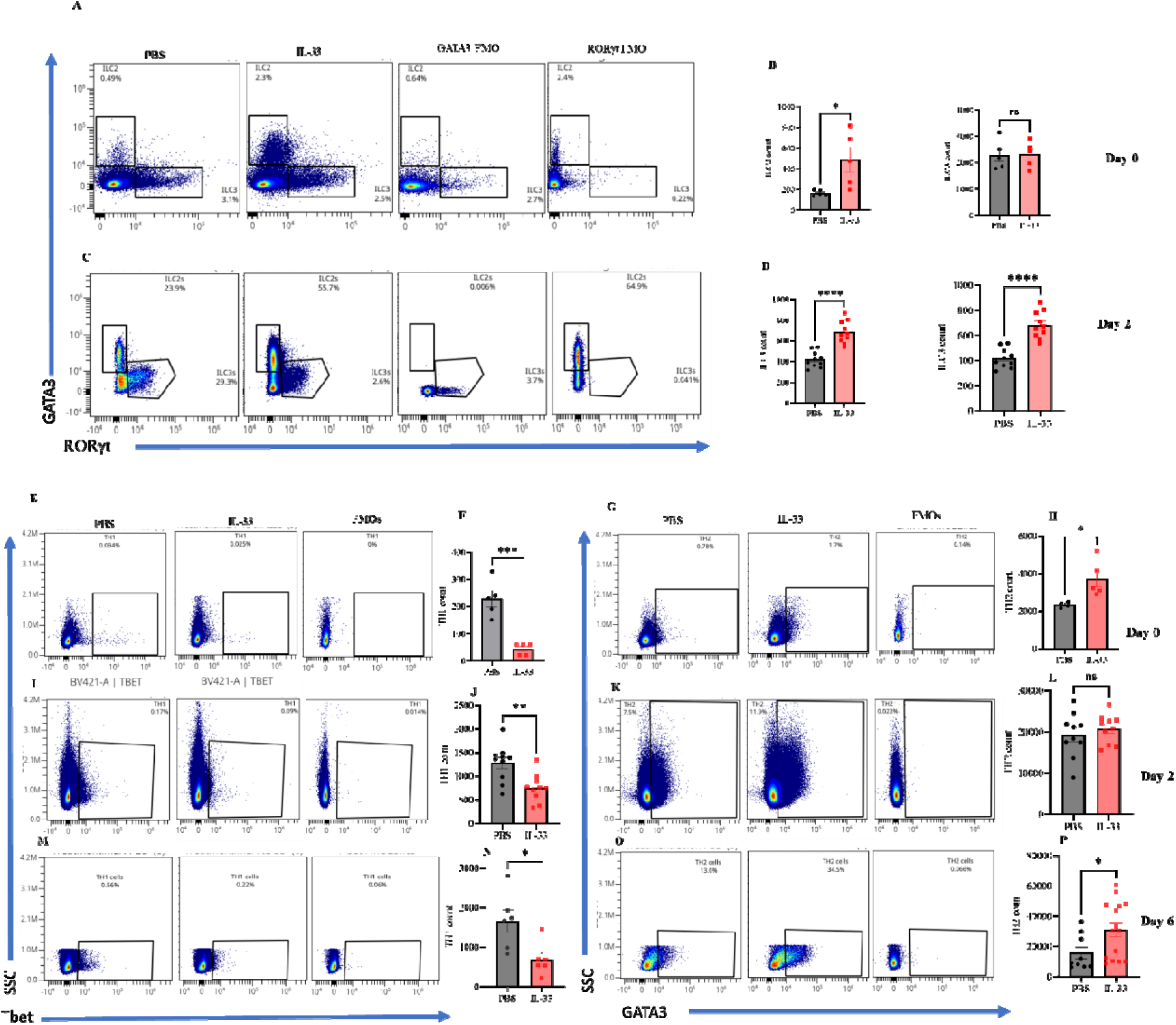
IL-33 increased mesenteric lymph node ILC2 and decreased TH1 cells during 1st C. difficile infection. IL-33 (0.75 µg) was administered i.p. on days -4 to 0 and mice infected on day 0. (A, B) ILC populations on day 0, prior to infection, and (C, D) at 2 days post-infection. TH1, TH2 populations (E-H) before infection, day 0; (I-L) day 2; and (M-P) day 6 post-infection. A two-tailed t-test for normally distributed data and a Mann-Whitney test for non-normally distributed data were used. Statistical significance is demarked as *P < 0.05, **P < 0.01, ***P < 0.001, and ****P < <0.0001. The error bar indicates SEM.

**Fig 6:**
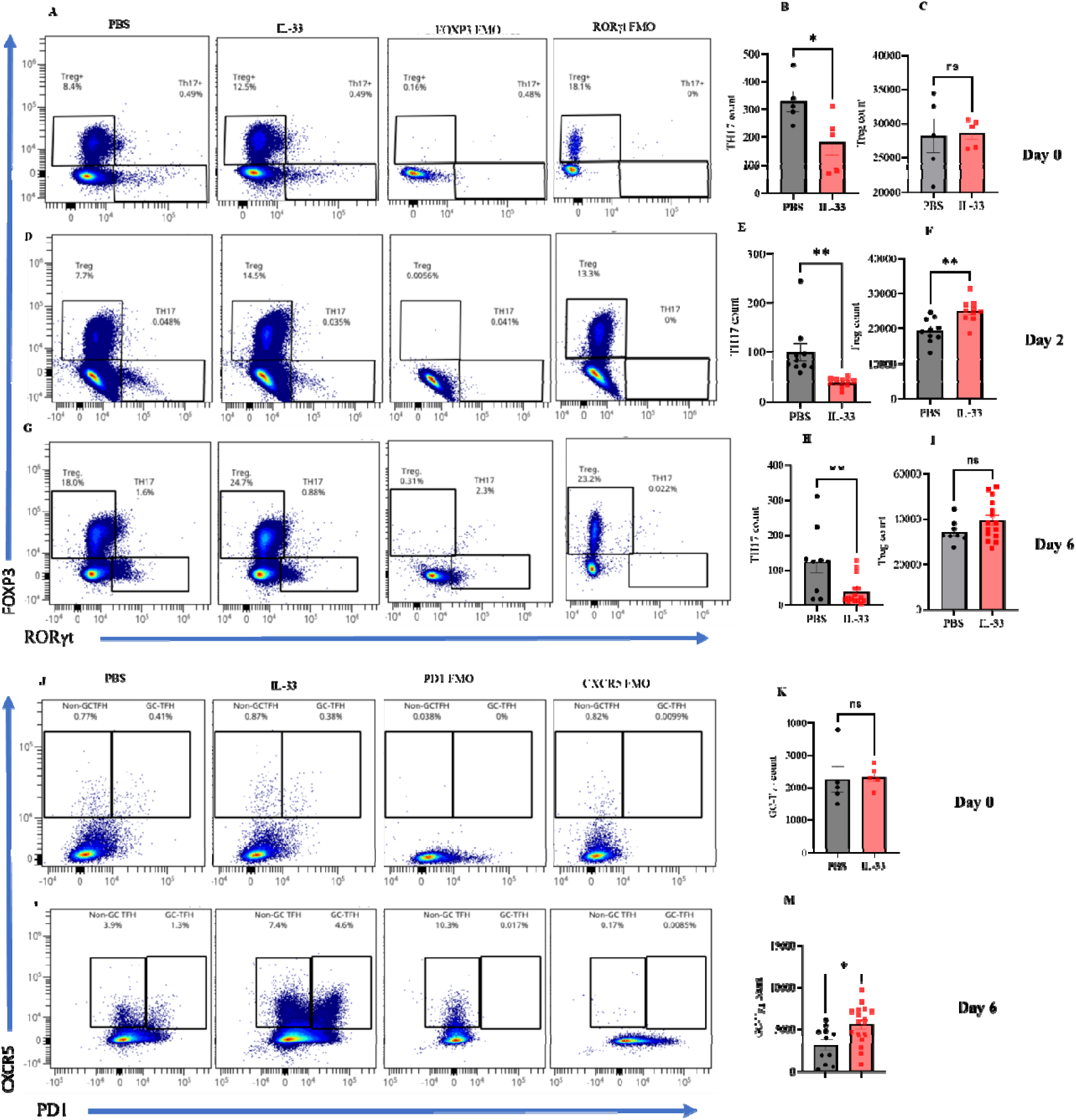
IL-33 increased mesenteric lymph node GC-TFH, Tregs, and decreased TH17 cells during a primary C. difficile infection. IL-33 (0.75 µg) was administered i.p. on days -4 to 0 and mice infected on day 0. Mesenteric lymph nodes were harvested to analyze T helper cells before infection, day 2, and day 6 post-first infection. (A-C) TH17 cells (CD45+, CD3+, CD4+, RORγt+) and Treg cells (CD45+, CD3+, CD4+, FOXP3+) on day 0 prior before infection; (D-F) on day 2; (G-I) on day 6 post-infection. (J-M) TFH cells were defined as germinal center (GC) TFH (CD45+ CD3+ CD4+ CD44+ PD1 high CXCR5+) and non-GC TFH cells (CD45+ CD3+ CD4+ CD44+ PD1 low CXCR5+) by flow cytometry. (J-K) TFH subsets on day 0 prior to infection; (L-M) TFH subsets on day 6 post-infection. A two-tailed t-test for normally distributed data and a Mann-Whitney test for non-normally distributed data were used. Statistical significance is demarked as *P < 0.05, **P < 0.01, and ***P < 0.001. The error bar indicates SEM.

### IL–33–mediated increase in activated mesenteric lymph node GC-TFH during primary *C. difficile* infection

CDI induces an inferior IgG response and is associated with a lack of T follicular helper cell (TFH) expansion (20). We hypothesized that IL-33 protection from recurrent CDI was due to TFH expansion to promote anti-toxin B antibody. We chose to measure Tfh cells on days 0 and 6 as this is consistent with the expected time it takes for Tfh cells to differentiate in the germinal center (21, 22). In support of this hypothesis, we found that IL-33 treatment increased activated GC-TFH by 1.3% to 4.6% of activated GC-TFH (**Figs. 6J, 6K, 6L, and 6M**).

Because IL-33 could activate dendritic cells via the ST2 receptor, we tested activation markers on dendritic cells before (day 0) and at 2, and 5 days of infection **(Supplemental Figs. 7A-7I).** There was a significant influx of CD11c-positive dendritic cells in MLN in the IL-33 treated group before infection but very less dendritic cells were found to have activation markers i.e. CD86 **(Supplemental Figs. 7A, 7B, 7C)**. At day 2 of infection, significantly more CD11c+ dendritic cells with significantly higher expressing activation marker CD86 were found in MLN **(Supplemental Figs. 7D, 7E, 7F)**, Which was not different by day 5 of infection **(Supplemental Figs. 7G, 7H, 7I)**. We concluded that IL-33 promoted anti-toxin B antibody by TFH expansion and in part by recruiting dendritic cells to the MLN.

**Fig 7:**
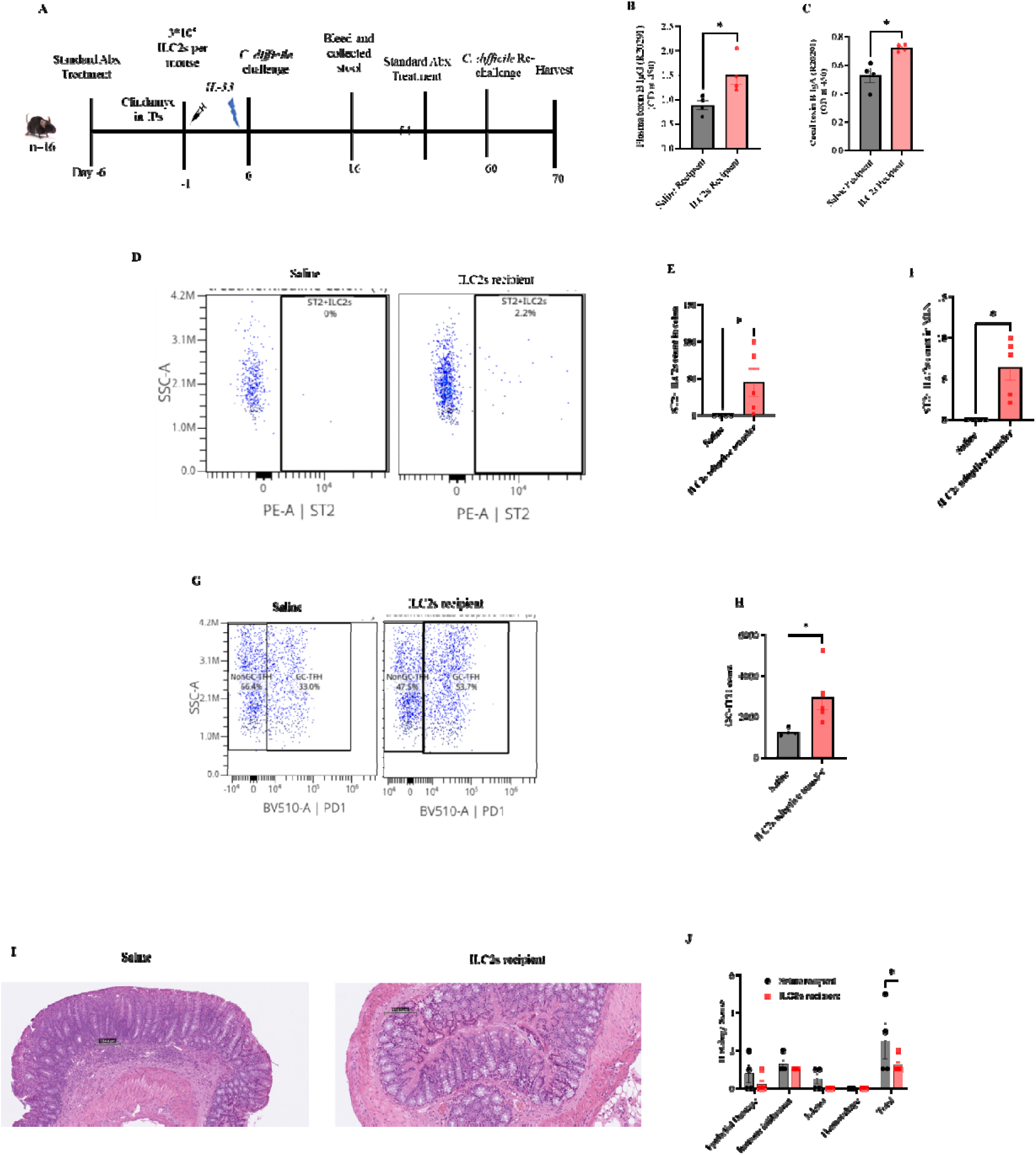
Adoptive transfer of ST2+ ILC2s into ST2 KO mice induces TFH cells and increased Toxin B specific antibody to protect from reinfection. ST2+ ILC2s (from uninfected IL-33 treated mice) were ex-vivo expanded, purified by flow-sorting and adoptively transferred into ST2 KO mice. **(A-C)** Mice were pretreated with antibiotics and injected with 0.75µg per dose per mouse of IL-33 in the gut one day after the adoptive transfer and one day before the R20291 infection. At 16 days post-primary infection, plasma toxin B-specific **(B)** IgG and **(C)** IgA antibodies were measured in plasma. **(D-N)** Mice were rechallenged with *C. difficile* on day 60. On day 70 (10 days post-2^nd^ infection), **(D-F)**, ILC2 were measured in the colon and mesenteric lymph nodes. **(G-H)** GC-TFH measured in the MLN **(I)** Day 70 representative epithelial damage (H&E) of treatment groups and **(J)** assessed by blinded scoring of infected tissue. A two-tailed t-test for normally distributed data and a Mann-Whitney test for non-normally distributed data were used. J, Šídák’s multiple comparisons test was used. Statistical significance is demarked as *P < 0.05, **P < 0.01, and ***P < 0.001. The error bar indicates SEM.

### Role of ILC2s in IL33-mediated protection from reinfection

We wanted to determine which primary or upstream cells responded to IL-33 via the ST2 receptor. We hypothesized that it was ILC2s that responded to IL-33 to promote anti-toxin B antibodies and protect from recurrence, in part because of their known role in adaptive immunity(23) and due to the observed increase in ILC2s in the IL-33 treated group after infection in MLN and colon **(Figs. 5A, 5B, 5C, and 5D & Supplemental Figs. 6A and 6B)**.

To test the role of ILC2 in IL-33-mediated anti-toxin B antibody production, ST2^+^ ILC2s were isolated from the spleen, mesenteric lymph nodes, and colon of IL-33-treated mice, expanded *in vitro*(24, 25), flow sort purified (**Supplemental Fig. 8)** and adoptively transferred into ST2^-/-^ mice (**Fig. 7A**). ST2^-/-^ mice that received ST2^+^ ILC2s had increased plasma anti-toxin B IgG and stool IgA following primary challenge with *C. difficile* (**Figs. 7B and 7C**). The presence of donor ST2+ ILC2s within the colon and MLN of recipient ST2^-/-^ mice was confirmed (**Figs. 7D, 7E, and 7F**). ST2^+^ ILC2 recipient mice had an increase in activated GC-TFH population (**Figs. 7G and 7H**). Upon reinfection, ST2^+^ ILC2 recipient mice had improved gut permeability, significantly more Goblet cells, and less epithelial damage (**Figs. 7I and 7J**). It is concluded that IL-33 protection from recurrent *C. difficile* was mediated by ILC2s **(Graphical abstract).**

### IL-33 is a biomarker for recurrent *C. difficile* infection

Utilizing a commercial multiplex proximity extension assay, IL-33 was measured in the blood of 58 hospitalized patients with CDI (within 48 hours of diagnosis) and 17 healthy controls. Patient details including demographics, comorbidities, etc. described in the supplementary table **(Supplementary Table. 1)**. Among the *C. difficile*-infected patients, 12 developed recurrent infections, and another 5 died within eight weeks. IL-33 was elevated in uncomplicated CDI (median 0.309 pg/mL) compared to healthy controls (median 0.068 pg/mL; Wilcoxon P<0.001). Only three out of the 45 cytokines measured significantly differed between patients who developed recurrent infection and those who did not. These cytokines included IL-33, C-X-C motif chemokine 10 (CXCL-10), and Tumor Necrosis Factor (TNF) (**Supplemental Figs. 10A). Figure 10B** presents ROC curve analysis for univariate and multivariable logistic regression models predicting recurrent *C. difficile* infection within 8 weeks. The univariate model, which includes only IL-33, performs similarly to the multivariable model that incorporates all three significant cytokines (IL-33, CXCL-10, and TNF). This suggests that while CXCL-10 and TNF are altered in patients with recurrent infection, they do not significantly improve the predictive power of the IL-33 model, which performs comparably in univariate and multivariable settings. IL-33 was higher in patients who went on to develop recurrent infection (median 0.600 pg/mL; P=0.031) compared to uncomplicated infection (**Figs. 8A and 8B**). Immunohistochemical staining revealed abundant anti-IL-33 staining of colonic epithelium from three non-recurrent and three recurrent human CDI patients (**Fig. 8C**).

**Fig 8:**
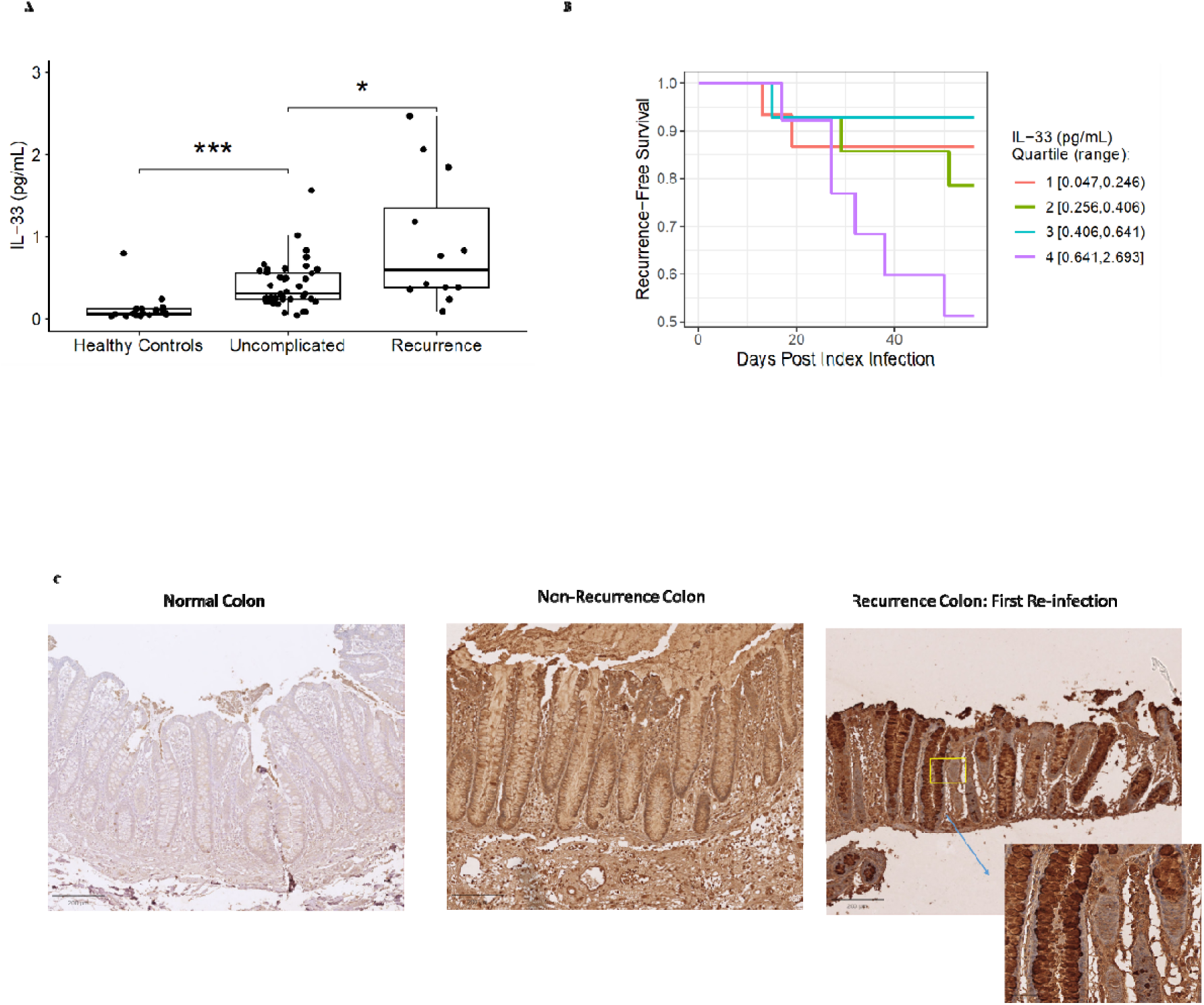
IL-33 is a biomarker for recurrent C. difficile infection in humans. (A) Plasma IL-33 was measured in healthy controls (n = 17), patients with uncomplicated CDI (n=41), and recurrent CDI (n=12) (excluding 5 patients who died) within 8 weeks of diagnosis. (B) Recurrence-free survival among the patients with *C. difficile* infection (1A), grouped by serum IL-33 quartile (Wilcoxon P=0.002). (C) Immunohistochemical staining of IL-33 from colon tissue biopsies of patients without or with recurrence. The Kaplan–Meier method was used to measure recurrence-free survival curves and to evaluate the effects of IL-33 on risk for recurrent C. difficile infection. To account for competing risk against recurrent infection, death within the 8-week follow-up period was treated as a censoring event.

### Discussion and Conclusion

*C. difficile* patients with the highest quartile serum IL-33 levels (>0.641 pg/mL) measured at diagnosis were more than 2.5 times more likely to develop recurrent infection within the following 8 weeks compared to patients with lower IL-33, suggesting IL-33 could serve as an early biomarker for reinfection comparing with the addition of CXCL-10 and TNF that do not significantly improve performance of a univariate IL-33 predictive model for recurrence. We employed a mouse model of reinfection to elucidate the role of IL-33 in rCDI. Our investigation revealed that IL-33 exerts a protective effect against CDI recurrence by activating group 2 innate lymphoid (ILC2) cells. ILC2 activation promoted humoral immunity against *C. difficile.* Transferring ST2+ ILC2s to ST2^-/-^ mice alone was enough to stimulate the production of toxin B-specific antibodies and promote the formation of T follicular helper (TFH) cell populations, consistent with earlier studies that ILC2 contributed to the development of T follicular helper (TFH) cells and antibody production (26).

The IL-33 signaling pathway plays a crucial role in promoting the humoral immune response and protecting against infection and reinfection by *C. difficile* colitis via the action of ILC2s. We observed that post-antibiotic dysbiosis, the administration of IL-33 resulted in elevated toxin B-specific antibody production, thereby conferring protection against toxin-induced epithelial damage and morbidity upon reinfection. To check whether the protection during reinfection is due to the IL-33-induced anti-toxinB antibodies, we utilized the µMT knock-out mouse model where B cells are depleted. Considering the limitations of this model as these mice could produce IgG and IgE antibodies(19), the αCD20 treatment mice model was used to reconfirm the results via depleteing *C. difficile* specific B cells. It is also important to note that αCD20 does not deplete terminally differentiated antibody-producing B220+ plasmablast and plasma cells that do not express CD20(27). These plasma cells could confer non-specific B cell antibody-mediated protection. Taking both results together, it was shown that IL-33-mediated protection during reinfection was due to toxin-specific antibodies. Our investigation corroborates earlier work showing that IL-33 facilitates the generation of IgA, contributing to the preservation of gut microbial homeostasis while mitigating IL-1α–induced colitis and colitis-associated cancer(28).

A likely target of humoral immunity is TcdB, which is recognized as the principal virulence determinant of *C. difficile.* TcdB has undergone expedited evolutionary changes, presumably in part to evade antibody neutralization. Clade 2 hypervirulent strains such as R20291 produce TcdB2, while VPI 10463 strains produce TcdB1(2). The disparities in the sequences of the two TcdB forms result in modifications to antigenic epitopes, diminishing cross-neutralization effectiveness by antibodies directed toward the C-terminal domain(29). Consistent with IL-33 acting to prevent recurrence by promotion of anti-TcdB antibodies, IL-33 administration did not provide cross-protection from a different TcdB type.

The interaction between antigen-stimulated B cells and the T follicular helper (TFH) cell subset determines the subsequent course of immune activity, i.e., antibody production. TFH cells play a pivotal role in supporting activated B cells through cognate interactions, characterized by antigen-specific binding, and by secreting a repertoire of functionally significant cytokines, thereby contributing to the coordination and enhancement of the humoral immune response (20). (GC-TFH) orchestrate B cells in germinal centers, facilitating somatic hypermutation and class-switch recombination, resulting in high-affinity antibodies. GC-TFH cells express specific molecules including CXCR5 and PD-1, enabling their precise modulation of B-cell interactions. Cytokines released by TFH augment B-cell differentiation into plasma cells, ultimately amplifying antibody secretion (30). Research has also shown that IL-33 has the potential to boost humoral immunity through its interactions with TFH cells(31). We found an IL-33-dependent increase in antibody production through increased GC-TFH activity.

There are several important limitations to this study. IL-33 was upregulated in patients at highest risk for recurrent CDI, suggesting that IL-33 could predispose to recurrent infection. However, our mouse data suggest that increased endogenous IL-33 is more likely a physiologic response to high-risk infection given the protective role of exogenous IL-33 in mice, stimulating protective humoral immunity. In addition, hospitalized patients with *C. difficile* infection were compared with healthy controls but were not adjusted for age, comorbidities, etc.

Here we have shown an additional role for IL-33 signaling in mitigating dysbiosis by downregulating TH17 and TH1 cells to create an environment inadequate for *C. difficile* disease. TH17 cells are crucial in elevating the risk of severe CDI by serving as a significant source of IL-17A(32). The sole adoptive transfer of TH17 cells has the potential to heighten the severity of CDI(32).

Understanding the role of IL-33 signaling in countering recurrent *C. difficile* infection (rCDI) through the ILC2-TFH axis is crucial for crafting potent CDI vaccines. A significant hurdle is addressing antigenic variation, notably in TcdB, demanding a nuanced approach for broad-spectrum defense. A successful vaccine must navigate these complexities to ensure comprehensive protection. IL-33-associated mechanisms present a promising avenue for advancing CDI vaccine strategies, fostering optimism for more effective preventive measures. A proposition arises to enhance vaccine effectiveness by supplementing the antigen with a type 2-skewed adjuvant, amplifying IL-33 signaling and fortifying protection against CDI recurrence.

In summary, our investigation reveals that IL-33 prompts the expansion of ILC2s, which in turn, either directly or indirectly, amplify TFH cells, supporting B cells in antibody production. The resulting toxin-specific antibodies are essential in mitigating clinical illness due to reinfection.

## Materials And Methods

### Mice

Our study exclusively examined male mice because the disease modeled is only relevant in males as females get less severe diseases. All the animal procedures were approved by the Institutional Animal Care and Use Committee at the University of Virginia (IACUC). C57BL/6J and μMT mice were purchased from the Jackson Laboratory, and ST2^-/-^ mice were obtained from Dr. Andrew McKenzie (Laboratory of Molecular Biology, Cambridge University, Cambridge, UK) (33). Sex-matched 8-12-week male mice were used in experiments. Animals were housed in a specific pathogen-free environment at the University of Virginia’s animal facility. The bedding was exchanged every 2 days for a minimum of 3 weeks to equilibrate their microbiota. Mice were infected with *C. difficile* as previously described (14). In short, mice were given an antibiotic cocktail of 215Dmg/L metronidazole (Hospira), 35Dmg/L colistin (Sigma), 45Dmg/L vancomycin (Mylan), and 35Dmg/L gentamicin (Sigma) in drinking water before 6 days of infection for 3 days. After 3 days, the mice were moved to regular drinking water. One day before the infection, clindamycin (Hospira) (0.016Dmg/g) was administered i.p. After infection, mice were monitored twice daily to evaluate clinical scoring parameters and weight loss over the course of infection and reinfection. As described elsewhere, the scoring criteria were weight loss, coat condition, eye condition, activity level, diarrhea, and posture (14). If mice reached a clinical score of 14 or lost more than 25% of weight, they were humanely euthanized. We define the acute phase as when the mice have active diarrhea and lose their weight and recovery is when the mice start gaining weight.

### Bacterial strains and culture

For the first infection, mice were infected with 5D×D10^4^ CFU/mL vegetative cells of either R20291 or VPI 10643 (ATCC 43244) strain. For reinfection, 10^6^ spores/ml of the R20291 strain or 5.2D×D10^4^ spores/ml of the VPI 10643 strain were used. *C. difficile* strains were plated on BHI agar from the glycerol stocks and incubated overnight at 37° C in an anaerobic chamber (34). Columbia, clospore, and BHI broth were reduced for at least 24 hours.

### To generate spore stocks

A single colony was inoculated into 15 ml of Columbia broth overnight at 37 °C, and then 5 ml of this culture was added to 45 ml of Clospore broth anaerobically and left for 7 days at 37 °C (35). The culture was washed with cold sterile water at least 5 times and resuspended in 1 ml of sterile water. The spores were stored in a 1.5 ml twist cap tube at 4 °C (Corning # 4309309).

### For vegetative infection

A single colony was inoculated in BHI medium overnight at 37 °C. The next day, cultures were washed twice with anaerobic PBS. The concentrations were measured by optical density for R20291 infection. For infection with the VPI 10643 (ATCC 43244) strain, the overnight culture was subcultured for 5 h before optical density measurement. The needed concentrations of vegetative cells were prepared and loaded into a syringe with a gavage needle inside the anaerobic chamber. Each mouse received 100 μl (5D×D10^3^ CFU for R20291 and 5D×D10^3^ CFU for VPI 10643) of inoculum by oral gavage. The actual inoculum was checked further by plating on BHI agar supplemented with 0.032 mg/mL cefoxitin, 1 mg/mL D-cycloserine, and 1% sodium taurocholate (Sigma) anaerobically incubated at 37 °C overnight. *C. difficile* burden was measured either by toxin A and toxin B specific qPCR on the DNA isolated from the stool or cecal content using a QIAamp fast DNA stool mini kit according to the manufacturer’s instructions or by using ELISA kit (TechLab Inc., catalog #TL5025) according to the manufacturer’s instructions to quantify bacterial count, normalized to stool or cecal content weight. The TOX A/B II ELISA kit (TechLab Inc., catalog #T5015) was used according to the manufacturer’s instructions to quantify Toxin A/B, normalized to stool or cecal content weight.

### Antibodies

Horseradish peroxidase (HRP)-conjugated anti-mouse IgM, IgG, IgG1, and IgA were purchased from Southern Biotech (Birmingham, AL), and goat anti-human IgG and Fcγ fragment specific antibodies from the Jackson Laboratory (109035-098).

### Tissue transcript and protein analysis

Tissue lysates were obtained by washing the ceca with 1X PBS and beating them for 1 minute in 300 µl of lysis buffer I containing 5 mM HEPES with 1X HALT protease inhibitor (Pierce). Next, tubes were incubated for 30 min on ice after adding 300 µl of buffer II containing 5 mM HEPES with 1X HALT protease inhibitor and 2% Triton X-100. The supernatant was collected after centrifugation at 13,000D×Dg at 4° C, and the total protein concentration was measured by BCA assay according to the manufacturer’s instructions (Pierce). The mouse Duoset sandwich ELISA kit (R&D) was used according to the manufacturer’s instructions to detect the IL-33 in the cecal tissue lysates.

For the IL-33 mRNA transcript analysis, the Rneasy mini kit from Qiagen and DNAse digestion ( TURBO DNA-free™ Kit from Invitrogen™) was used according to the manufacturer’s instructions. RNA from the cecal tissue was stored in RNA-later at -80 °C. Tetro cDNA synthesis kit (Bioline) was used to prepare the cDNA, and amplification of IL-33 was done by Taqman IL-33 primer/probe set (Applied Biosciences: Mm00505403_m1). Normalization of gene expression was done by HPRT and GAPDH housekeeping genes.

### Flow cytometry

The colon was rinsed in a buffer containing HBSS, 25 mM HEPES, and 5% FBS. Dissociation buffer (HBSS with 15 mM HEPES, 5 mM EDTA, 10% FBS, and 1 mM DTT) was used for 40 minutes at 37 ° C with 122 rpm agitation to remove the epithelial cells from the isolated colon. Digestion buffer (RPMI 1640 containing 0.17 mg/mL Liberase TL (Roche) and 30 µg/mL DNase (Sigma)) was used for 40 minutes at 37 ° C with 122 rpm agitation to digest manually diced lamina propria. After digestion, a 100 µM cell strainer followed by a 40 µM cell strainer (Fisher Scientific) was used to get single-cell suspensions. Extracellular staining was done with BB515-CD19 (BD 564509, dilution 1/25), PerCP-Cy5.5-CD5 (100624, dilution 1/100), PerCP-Cy5.5-CD3 (100218, dilution 1/100), PerCP-Cy5.5-FcεRIα (134320, dilution 1/100), BV510-CD90 (140319, dilution 1/25), BUV805-CD11b (368-011282, dilution 1/400), PE-CY5-CD64 (139332, dilution 1/75), APC-Fire 810-Ly6C (128055, dilution 1/400), BV785-CD45 (103149, dilution 1/200), NovaFluor Blue 610-70s-CD8a (M003T02B06, dilution 1/200), AF700-CD4 (100430, dilution 1/400), FITC-TCRbeta (159706, dilution 1/50), BV605-TCRgd (118129, dilution 1/200), BV650-CD11c(117339, dilution 1/100), APC Cy7-CD103 (121432, dilution 1/30), Pacific blue-CD40 (124626, dilution 1/50), AF647-CD80 (104718, dilution 1/100), PE-CD86 (105008, dilution 1/50), BUV 737-Ly6G (367-9668-82, dilution 1.25/100), Sprk UV 387-MHCII (107670. Dilution 1/400), BUV 563-SiglecF (365-1702-82, dilution 1/400), PE-Dazzle 594-CD127 (135032, dilution 1/100), PE Dazzle 594-CXCR5 (145522, dilution 1/100), FITC-CD44 (103005, dilution 1/100), BV510-PD1 (135241, dilution 1/100). Intracellular staining was done with BV421-Tbet (5563318, dilution 1/20), APC-RoryT (1769818, dilution 1/33), BV711-GATA3 (565449, dilution 1/50), PerCP-eFluor 710-FOXP3(46577382, dilution 1/100), and PE-fire 700-CD206 (141741, dilution 1/75). For surface staining, 1D×D10^6^ cells/sample were Fc-blocked with TruStain fcX (BioLegend, #101320, 1/200) for ten minutes at room temperature followed by LIVE/DEAD blue (Thermoscientific L34962) for 30 min at 4 °C. Cells were washed twice in FACS buffer (PBS+ 2% FBS) and stained with fluorochrome-conjugated antibodies for 30 min at 4° C. Cells were washed and resuspended in Foxp3 Fix/Perm WorkinFg Solution (ebiosciences, #00-5523-00) and incubated overnight at 4 °C. Cells were washed twice with permeabilization buffer and stained for 30 min at room temperature. Flow cytometry was performed on a Cytek Aurora (5-Laser) Spectral Flow Cytometer and analysis was done on Omiq software. All cell counts were normalized based on 80,000 live cell counts. SpectroFlo QC beads (SKU B7-10001) were used for routine performance tracking of the Cytek Aurora (5-Laser) Spectral Flow Cytometer. Unmixing was done using single stains prepared on either cells or UltraComp eBeads™ Plus Compensation Beads (01-3333-42). All the gating strategies used in the article are presented in the supplementary figure **(Supplemental Fig. 10).**

### IL-33 and **α**CD20 Treatment

Carrier-free recombinant mouse IL-33 (Biolegend; Catalog #: 580504) was diluted with sterile PBS to prepare a 7.5 µg/ml solution. 100 μl was injected intraperitoneally daily for 5 days prior to the first infection or reinfection. αCD20 was a generous gift from Genentech. 250 µg of αCD20 per mouse was injected intraperitoneally at day -7, 3, 18, 33, 48, 63.

### ELISA

96 well half-area assay plates (Corning) were used to detect toxin-B-specific antibodies. Plates were coated with 2 μg/ ml of toxin B in carbonate buffer (Sigma), a generous gift from Techlab, and kept at 4 °C overnight. The next day, plates were blocked with 1% bovine serum albumin (BSA) in PBS and 0.05% Tween 20 at 37 °C for 1 hour. Samples were added by diluting mouse sera, fecal supernatant, or cecal content in PBS-T, and plates were kept at 37°C for another 1 hour. *C. difficile* toxin B monoclonal antibody, clone: A13I, from Invitrogen, was used as a positive control. Horseradish peroxidase (HRP)- conjugated IgG (1:5000), IgG1 (1:5000), IgM (1:1,000), or IgA (1:1000) was added after washing three times with PBS-T. Wells were developed by either 2,2’-azinobis(3-ethylbenzthiazolinesulfonic acid) (ABTS) substrate (KPL, Gaithersburg, MD) or ultra TMB-ELISA substrate solution (Thermo Scientific) at room temperature for 10 minutes and stopped by 1% SDS or 2 M H2SO4 respectively. Optical density was read at 410 nm, background at 650 nm for ABTS substrate, and 450 nm for TMB substrate. A well without a sample and control plasma was used as the negative control.

### ILC2 Adoptive Transfer Studies

Mesenteric lymph nodes (MLN), spleen, colon, and cecum were extracted from wild-type C57BL/6J mice that had received 5 daily doses of IL-33 (0.75 µg). A single-cell suspension was prepared from the colon and cecum, as described above. For MLN and spleen, a 40 µM cell strainer (Fisher Scientific) was used to prepare a single-cell suspension. After passing through the 40 µM cell strainer, 1 ml of 1X red blood cell (RBC) lysis buffer (Thermo Scientific) was added for 1 minute. After centrifugation at 1600 rpm for 6 minutes, single cells were prepared in FACS buffer containing 2% heat-inactivated fetal bovine serum (FBS) in PBS. The single cells obtained from MLN, spleen, cecal, and colon were subjected to lineage^+^ cell depletion by magnetic bead purification of lineage-negative populations (bulk ILCs) (Miltenyi Lineage Cell Depletion Kit: 130-110-470). The lineage-negative cells were expanded *in vitro* in complete RPMID1640 media containing 10% FBS, 2 mM glutamine, 100 U/ml penicillin, 100 μg/ml streptomycin, 50 ng/ml of IL-33, and 10 ng/ml of IL-2 and IL-7 for 4 days. Cells were flow sorted on the Influx Cell Sorter (BD Biosciences) based on Lin^-^ (CD11c, CD3, CD5, CD11b, CD19, Fc epsilon R1 alpha with PECy7 fluorochrome) CD45+ CD90.1+ CD127+ CD25+ ST2+ expression after cell surface staining. Approximately 3 × 10^5^ ILC2s were transferred into each ST2^-/-^ mouse.

### Detection of Human IL-33 in serum

Hospitalized patients between 18-90 years old with diarrhea and a positive CDI PCR test (nucleic acid amplification test (NAAT) GeneXpertSerum) were approached to be in the study. Patient demographics and clinical data were collected, and a follow-up was completed on all subjects to determine CDI recurrence and mortality after enrollment. Consented and enrolled subjects had 20 ml of blood drawn in EDTA tubes within 24 hours of CDI diagnosis. Blood was spun down at 2000xg for 15 minutes at room temperature and plasma was stored in aliquots at -80°C until utilized, at which point they were thawed on ice. Plasma were collected from 17 healthy donors without *C. difficile* infection, and 58 prospectively enrolled hospitalized patients within 48 hours of diagnosing *C. difficile* infection. Among *C. difficile*- infected patients, clinical outcomes (recurrent infection or death) were measured over an 8-week follow-up period. Recurrent *C. difficile* infection was defined as symptom relapse following completion of treatment for the index episode, requiring re-treatment. Serum cytokine concentrations were measured using a commercial multiplex proximity extension assay (Olink Proteomics; Watertown, MA). Concentrations of IL-33 were compared between healthy controls, patients who survived the follow-up period without recurrent *C. difficile* infection, versus those who developed a complication (recurrence or death). The collection and analyses of patient samples and healthy controls were approved by the University of Virginia Institutional Review Board (IBR-HSR18782 and HSR220013).

### Mouse and human histology and immunohistochemistry

Proximal colonic sections were fixed in Bouin’s solution and transferred to 70% ethanol after 24 h. Staining was done with either hematoxylin and eosin (H&E) or Periodic Acid Schiff (PAS) after preparing paraffin-embedded sections by the University of Virginia Research Histology Core. Two blinded observers scored histopathology using a scale from 0 to 3 for submucosal edema, We utilized: 0= none, 1= Mild, 2= Moderate, and 3= Intense/Severe damage for epithelial disruption and immune cell infiltration, and hemorrhage was scored as yes=1, No=0(36). Goblet cells were identified as PAS^+^ and their number normalized to the number of crypts. Human biopsies sourced from the University of Virginia Biorepository and Tissue Research Facility were utilized. Researchers were kept blind to patient identities. The University of Virginia Biorepository Core conducted staining of human colon biopsy sections using a primary antibody targeting IL-33 (R&D, AF3625, diluted at 1/80,000).

### FITC-Dextran Gut Permeability Assay

To assess intestinal permeability, mice were orally administered a fluorescein isothiocyanate (FITC)- dextran solution (Sigma-Aldrich, #46944-500MG-F) at a dosage of 44 mg per 100 g body weight. Four hours post-administration, mice were euthanized, and serum samples were collected. The concentration of FITC-dextran in the serum was measured using a spectrophotometer set at excitation and emission wavelengths of 485 nm and 530 nm, respectively.

### Statistical analyses

The Kaplan–Meier method was used to measure recurrence-free survival curves and to evaluate the effects of IL-33 on risk for recurrent *C. difficile* infection. To account for competing risk against recurrent infection, death within the 8-week follow-up period was treated as a censoring event. A two-tailed t-test for normally distributed data and a Mann-Whitney test for non-normally distributed data (serum IL-33) were used to determine the statistical significance between groups. Statistical analyses were performed using GraphPad Prism software (GraphPad Software Inc., La Jolla, CA) or R version 4.2.0 (R Core Team, Vienna, Austria).

## Acknowledgments

The authors thank the flow cytometry and research histology cores at the University of Virginia for providing their expertise. The authors thank TechLab, Inc. for generously sharing toxin B for ELISA and Toxin A/B reagents. The authors acknowledge Dr. Andrew McKenzie (Laboratory of Molecular Biology, Cambridge University, Cambridge, UK) for sharing ST2-deficient mice. The authors would like to acknowledge Genentech for providing αCD20 antibody. This work was supported by grants from the US National Institutes of Health (R01 AI152477 and R01 AI124214) to WAP.

## Author Contributions

FN and W. A. P. designed all of the experiments. F. N performed the experiments, analyzed and interpreted data, and wrote the manuscript. J. U. helped with tissue extraction and processing. R. H. B. and AB helped with mouse tissue extraction. D.T. helped in tissue biopsies scoring. MKY, G. R.; I. R.; and G. R. M. helped in human studies. G. R. M., G. R., D.T., and W. A. P. edited the manuscript. W. A. P. supported all aspects of the work.

## Resource Availability

Further information and requests for reagents will be fulfilled by the corresponding author William Petri or Farha Naz.

---

**Supplementary Fig 1:**
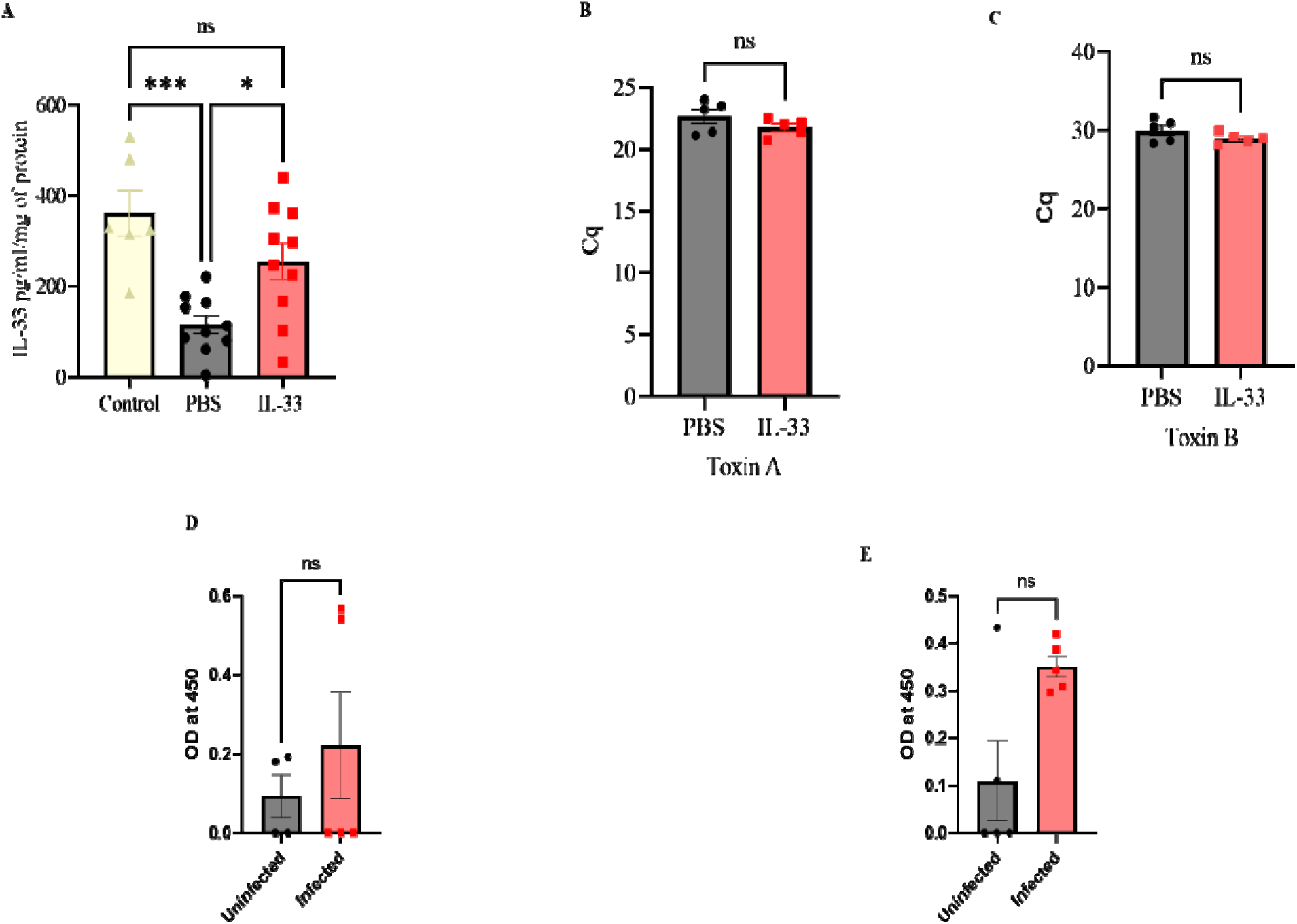
IL-33 treatment restores depleted IL-33 levels caused by antibiotics without altering C. difficile burden. In a mice model of CDI, IL-33 (0.75 µg) was administered i.e. on days -4 to 0 to C57BL/6J mice. (A) Proteins were isolated from the cecal tissue and IL-33 concentrations were determined with ELISA prior to infection; (B-C) DNA was isolated from cecal content after 15 days of infection, and qPCR was performed to check the bacterial load based on (B) Toxin A; and (C) for Toxin B. ST2−/− mice produces nonsignificant toxin-specific antibodies after first infection with C. difficile without IL-33 administration (D-E). On post-infection day 15, antibodies were measured in plasma. (D) plasma toxin B specific IgG from ST2 -/- mice uninfected and infected with C. difficile (E) plasma toxin B specific IgM from ST2 -/- mice uninfected and infected with C. difficile strain. A two-tailed t-test was used. Statistical significance is demarked as ns (non-significant), *P < 0.05, **P < 0.01, and ***P < 0.001. The error bar indicate SEM.

**Supplementary Fig 2:**
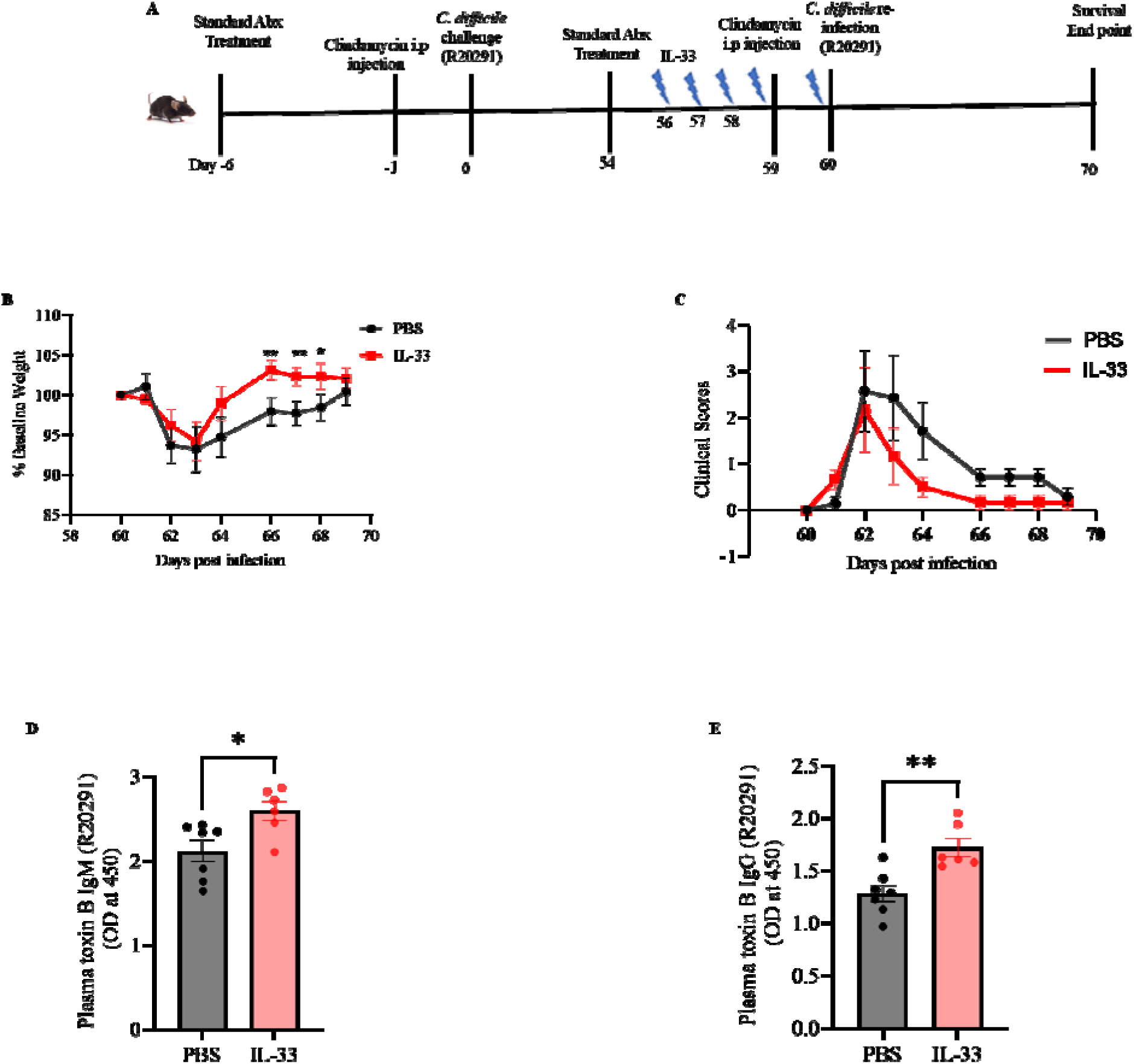
IL33 decreases the severity of a 2nd C. difficile infection when given after the first infection. IL-33 (0.75 µg) was administered i.p. on days 56 to 60 after the first infection and infected again on day 60 with the same C. difficile strain R20291. (A) Experimental design for IL-33 treatment and 2nd infection; (B) 2nd infection weight loss; (C) clinical scores; (D) plasma toxin B specific IgM; (E) plasma toxin B specific IgG. Comparison made by two-tailed Student’s t-test. *P < 0.05, **P < 0.01, and ***P < 0.001. The error bar indicates SEM.

**Supplementary Fig 3:**
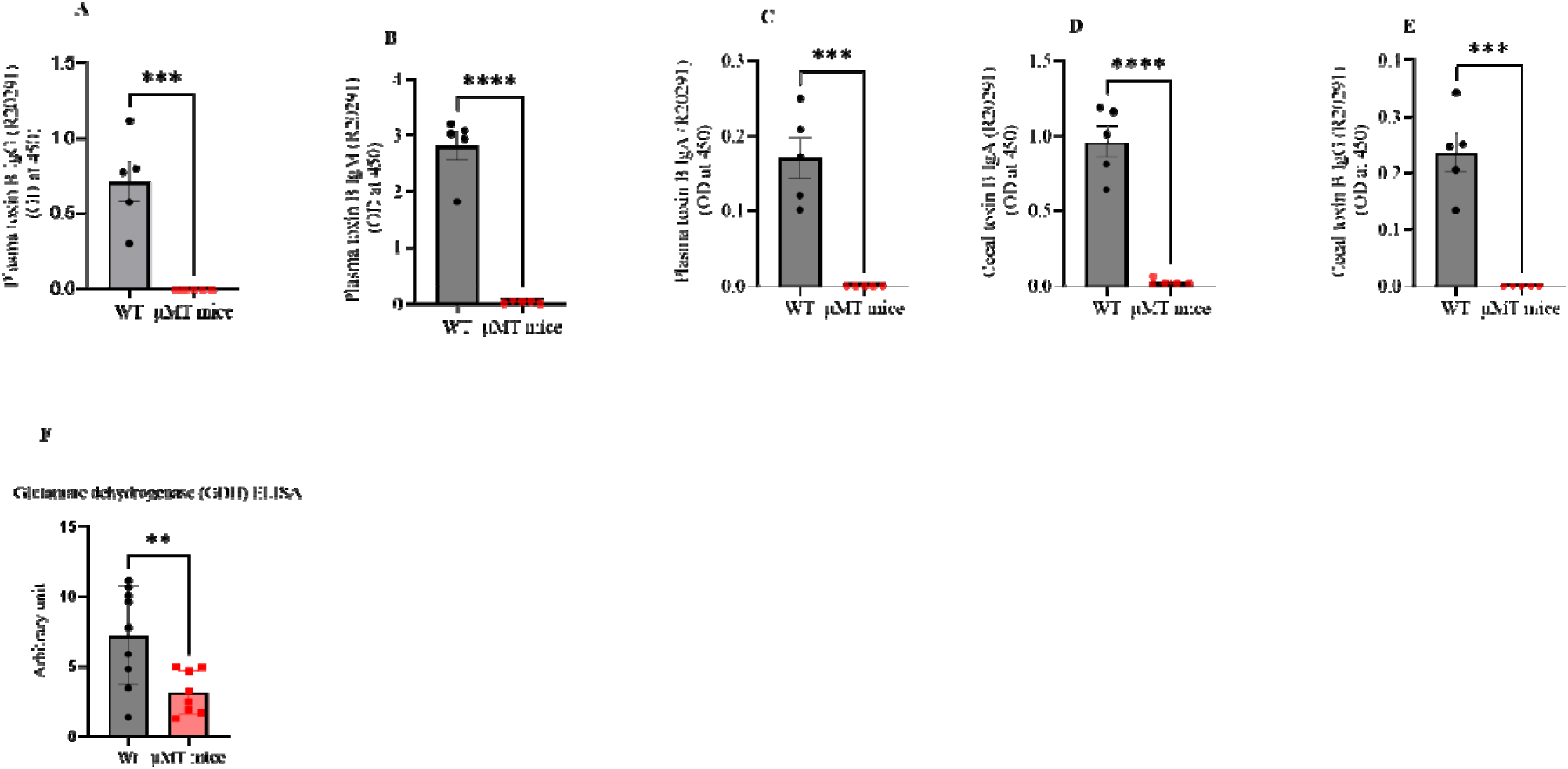
**μ**MT KO mice are not able to produce toxin-specific antibodies. WT and μMT KO mice were administered IL-33 (0.75 µg) i.p. on days -4 to 0 and mice infected on day 0 with C. difficile strain R20291. (A) Toxin B specific plasma IgG; (B) IgM; (C) IgA; (D) stool IgA; and (E) stool IgG measured on day 23 post-first infection. WT and μMT KO mice were reinfected with C. difficile R20291 60 days after primary infection. (F) Stool C. difficile burden measured by glutamate dehydrogenase ELISA. Comparison made by two-tailed Student’s t-test. *P < 0.05, **P < 0.01, and ***P < 0.001. The error bar indicates SEM.

**Supplementary Fig 4:**
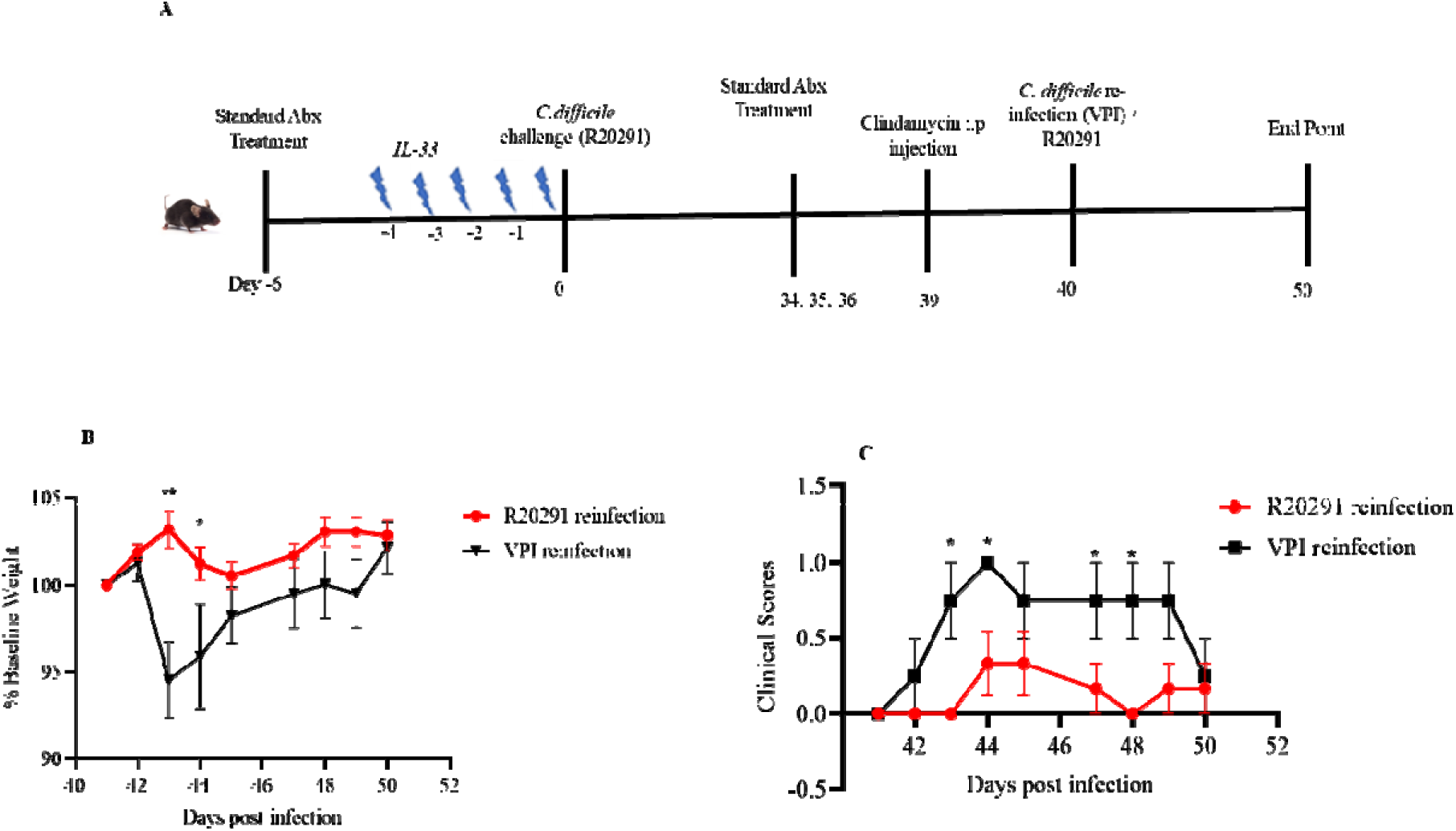
Lack of cross-protection by IL-33 with a first infection with TcdB2 R20291 strain and second with TcdB1 VPI strain. Mice were administered IL-33 (0.75 µg) i.p. on days -4 to 0 and infected on day 0 with C. difficile strain R20291 and again on day 40 with either C. difficile strain R20291 or the VPI strain. (A) Experimental design; (B) reinfection weight loss; (C) clinical scores. Comparison made by two-tailed Student’s t-test. Statistical significance is demarked as *P < 0.05, **P < 0.01, and ***P < 0.001. The error bar indicate SEM.

**Supplementary Fig 5:**
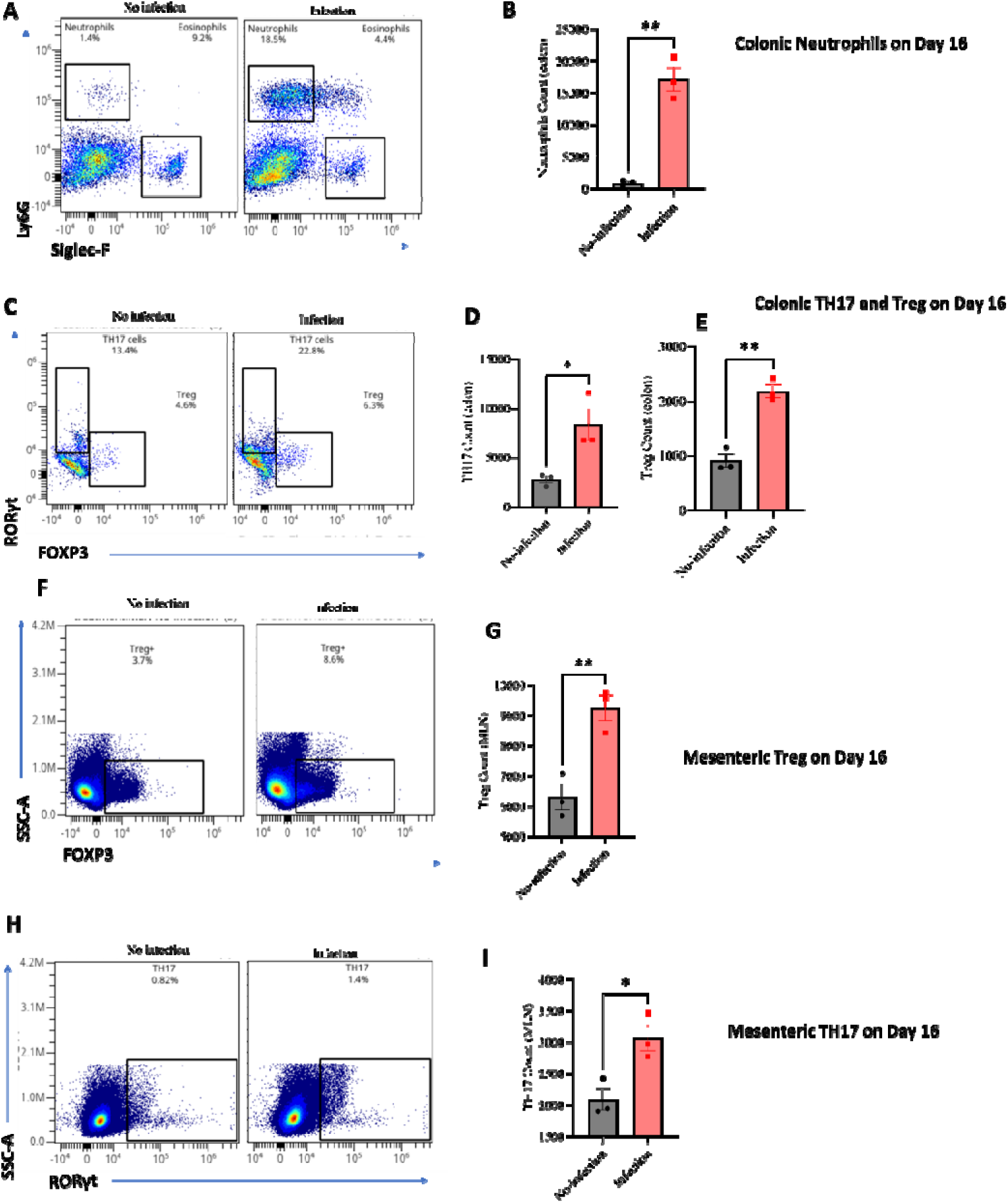
Th17 cells persist beyond the resolution of the disease. Mesenteric lymph nodes and colon were harvested for analysis of neutrophils and T helper cells on day 16 post-first infection. Colonic (A-B) neutrophils; (C-E) TH17 (CD45+, CD3+, CD4+, RORγt+) and Tregs (CD45+, CD3+, CD4+, FOXP3+). MLN (F-G) Treg and (H-I) TH17 cells. Comparison made by two-tailed Student’s t-test. Statistical significance is demarked as *P < 0.05, **P < 0.01, and ***P < 0.001. The error bar indicates SEM.

**Supplementary Fig 6:**
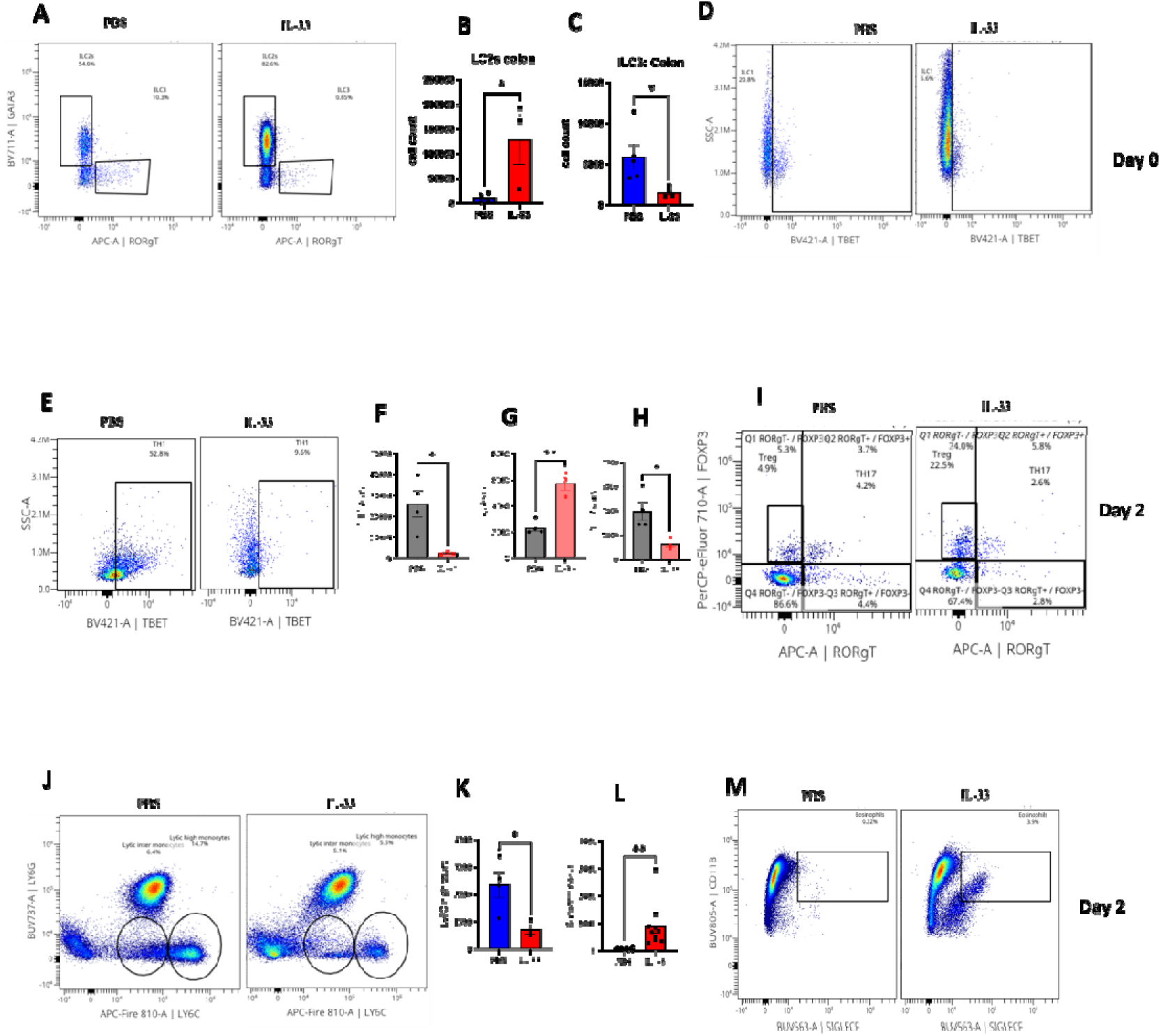
IL-33 increases colonic eosinophils, ILC2, Treg and decreased inflammatory monocytes, TH1, and TH17 cells during 1st C. difficile infection. IL-33 (0.75 µg) was administered i.p. on days -4 to 0 and mice infected on day 0. Innate lymphoid cells (ILC), myeloid cells, and T cells from colonic lamina propria were evaluated on days 0 and 2 post-first infection. (A-D) ILCs; (E-F) TH1 cells; (G, I) Treg and TH17 cells on day 0 prior to infection. (J-K) inflammatory monocytes; and (L-M) Eosinophils day 2 post-infection. Comparison made by two-tailed Student’s t-test. Statistical significance is demarked as *P < 0.05, **P < 0.01, and ***P < 0.001. Th error bar indicates SEM.

**Supplementary Fig 7:**
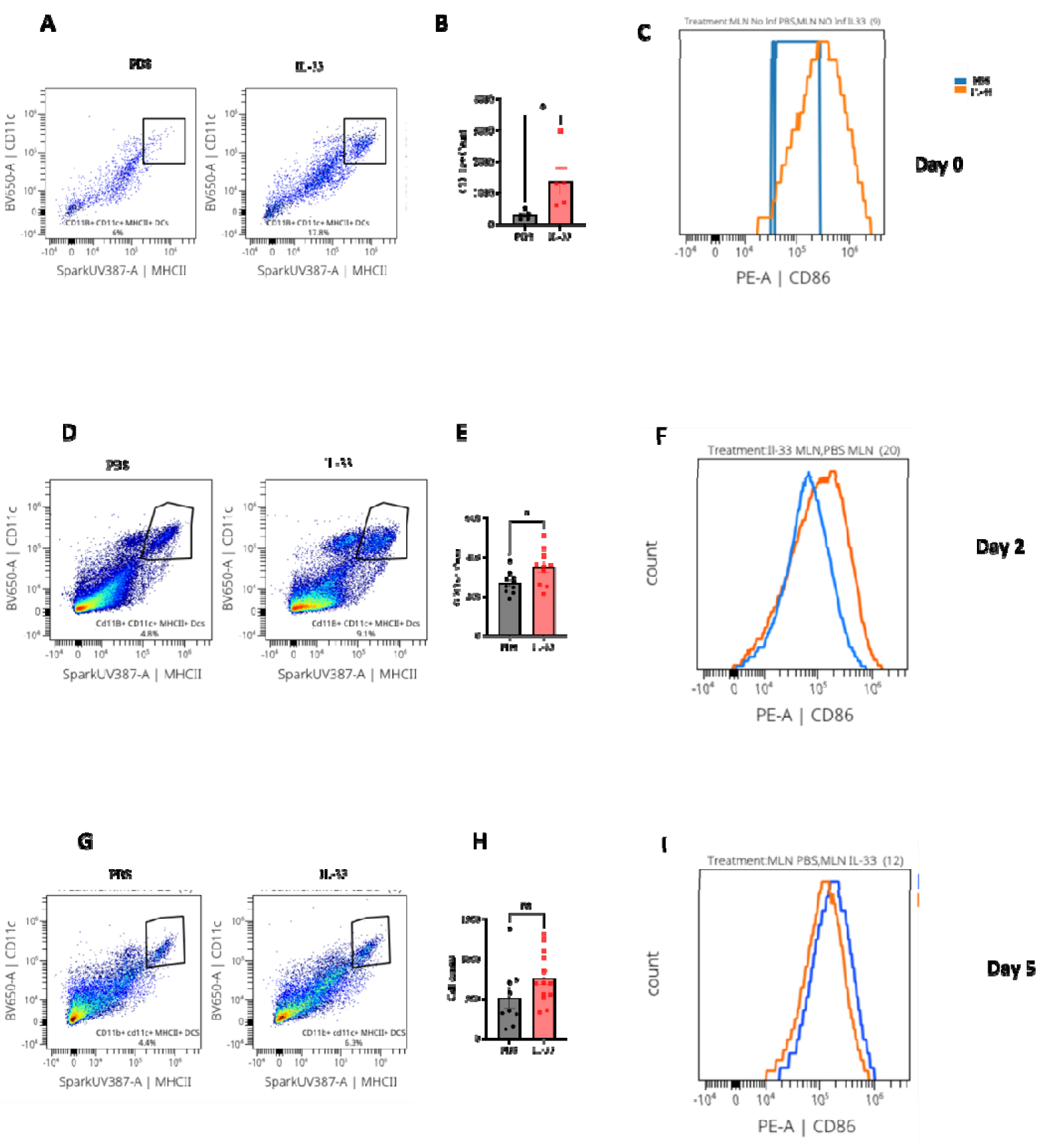
IL-33 increases the number of dendritic cells but did not increase their activation during a primary C. difficile infection. IL-33 (0.75 µg) was administered i.p. on days -4 to 0 and mice infected on day 0. Mesenteric lymph nodes were harvested for analysis of dendritic cells by flow cytometry before infection, day 2, and day 5 post-first infection. Dendritic cells (CD45+ CD11b+ MHCII+ CD11C+) (A-C) on day 0 prior before infection; (D-F) on day 2; (G-I) on day 5 post-infection and MFI of activation markers, CD86, CD80, and CD40 on dendritic cells (C, F, I) were analyzed. Comparison made by two-tailed Student’s t-test. Statistical significance is demarked as *P < 0.05, **P < 0.01, and ***P < 0.001. The error bar indicate SEM.

**Supplementary Fig 8:**
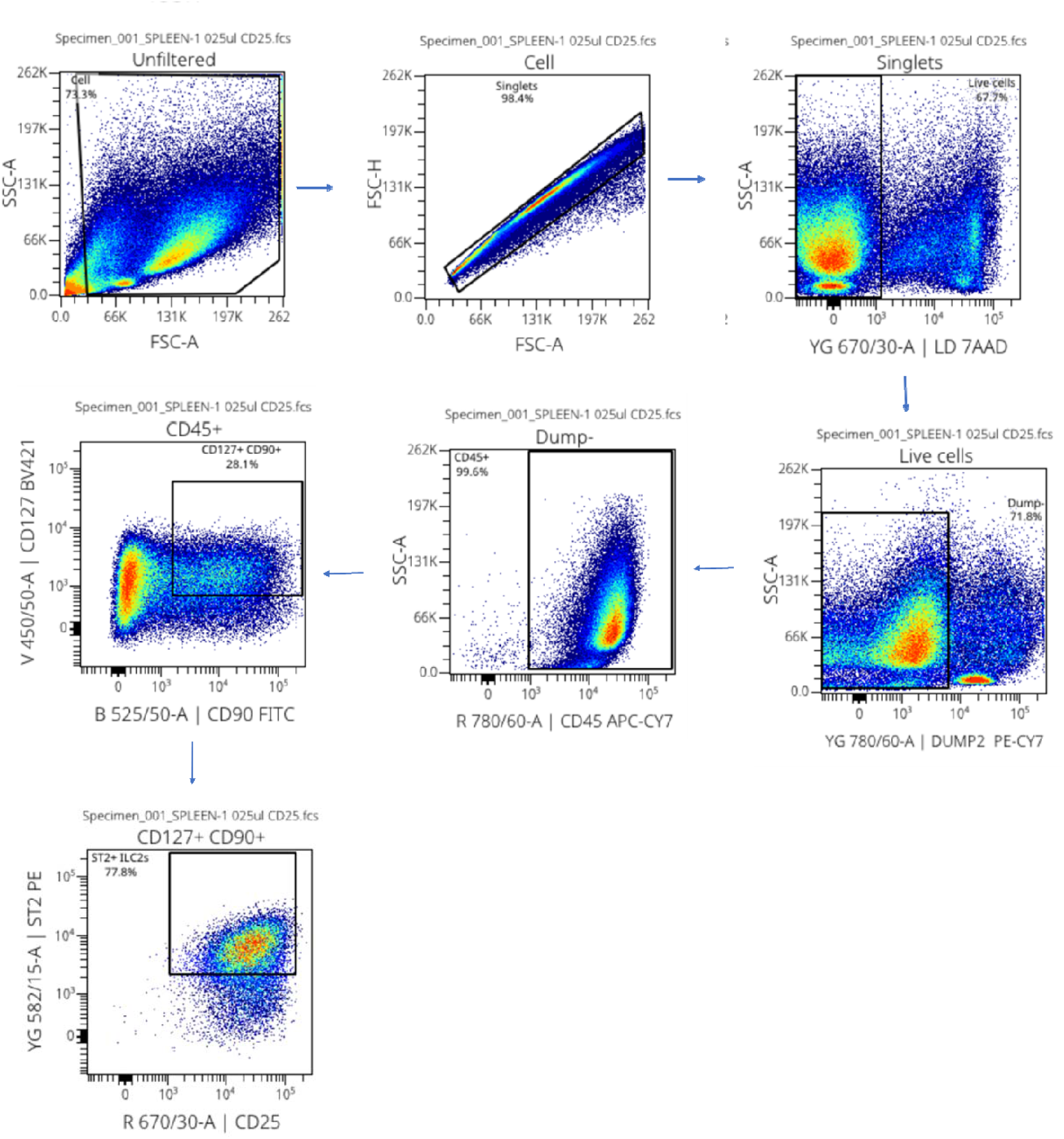
Gating strategy used for ST2+ ILC2s flow sorting for adoptive transfer.

**Supplementary Fig 9:**
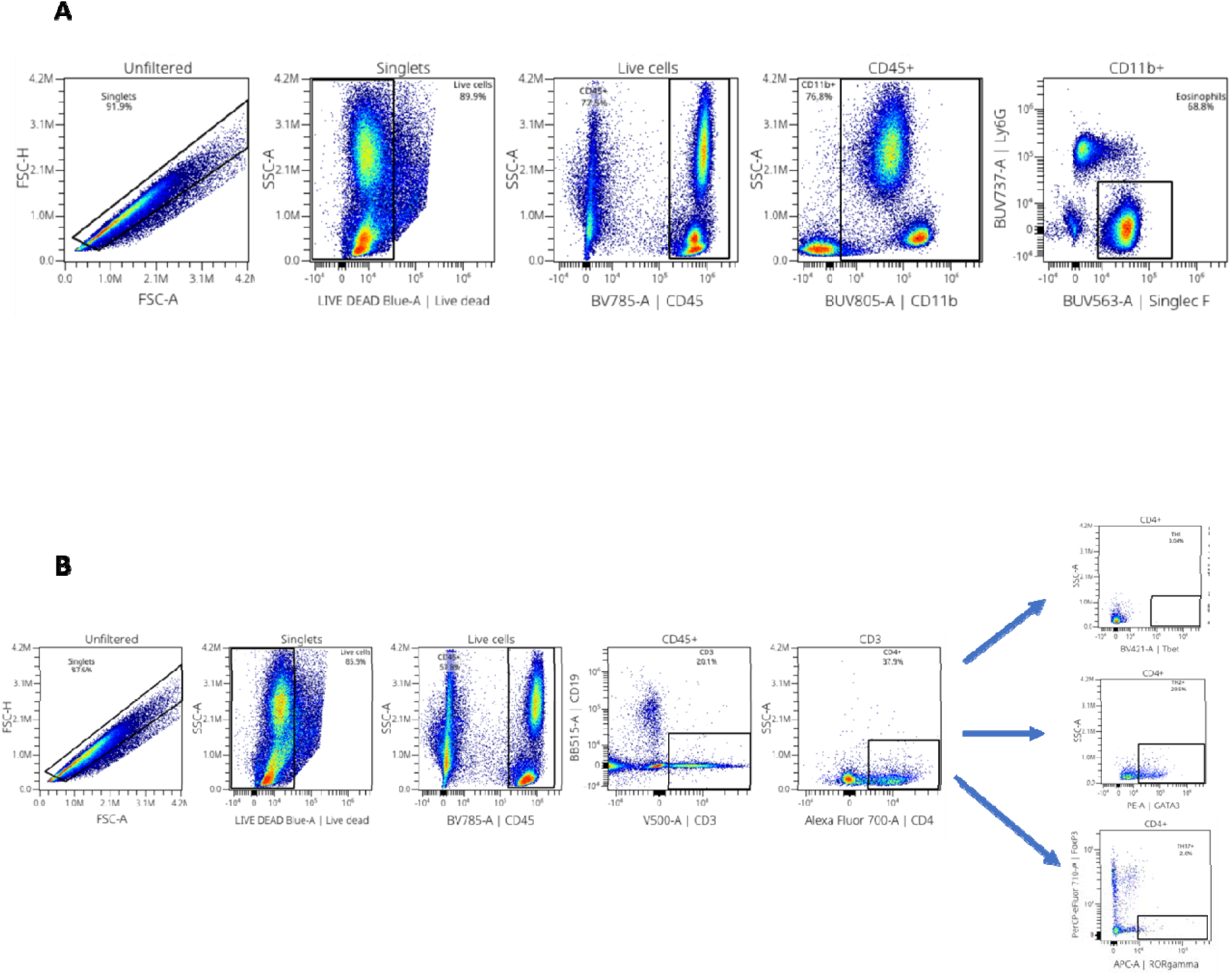
Gating strategy used for (A) Eosinophils and neutrophils; (B) TH1, TH2, TH17, and Treg cells.

**Supplementary Fig 10:**
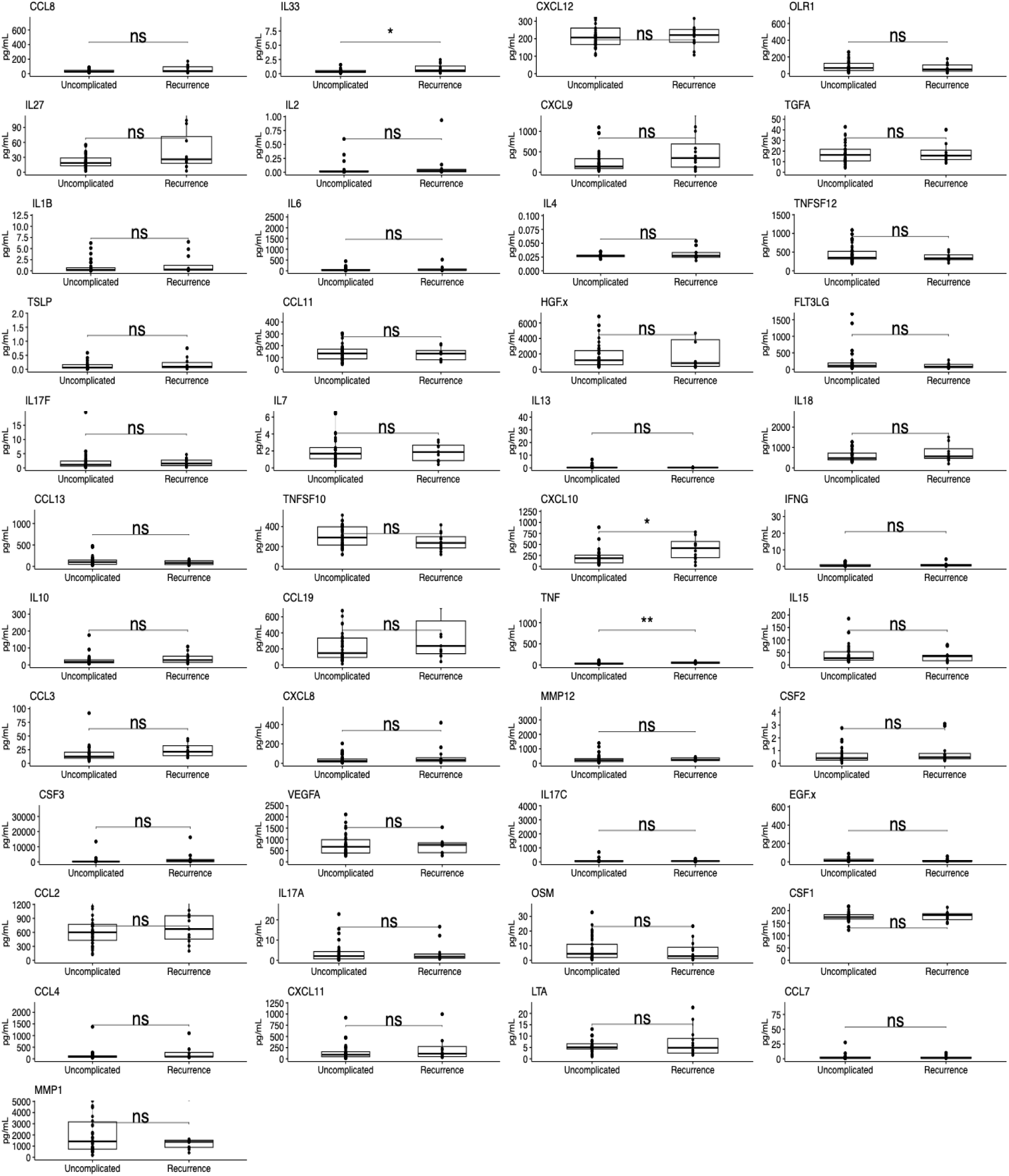
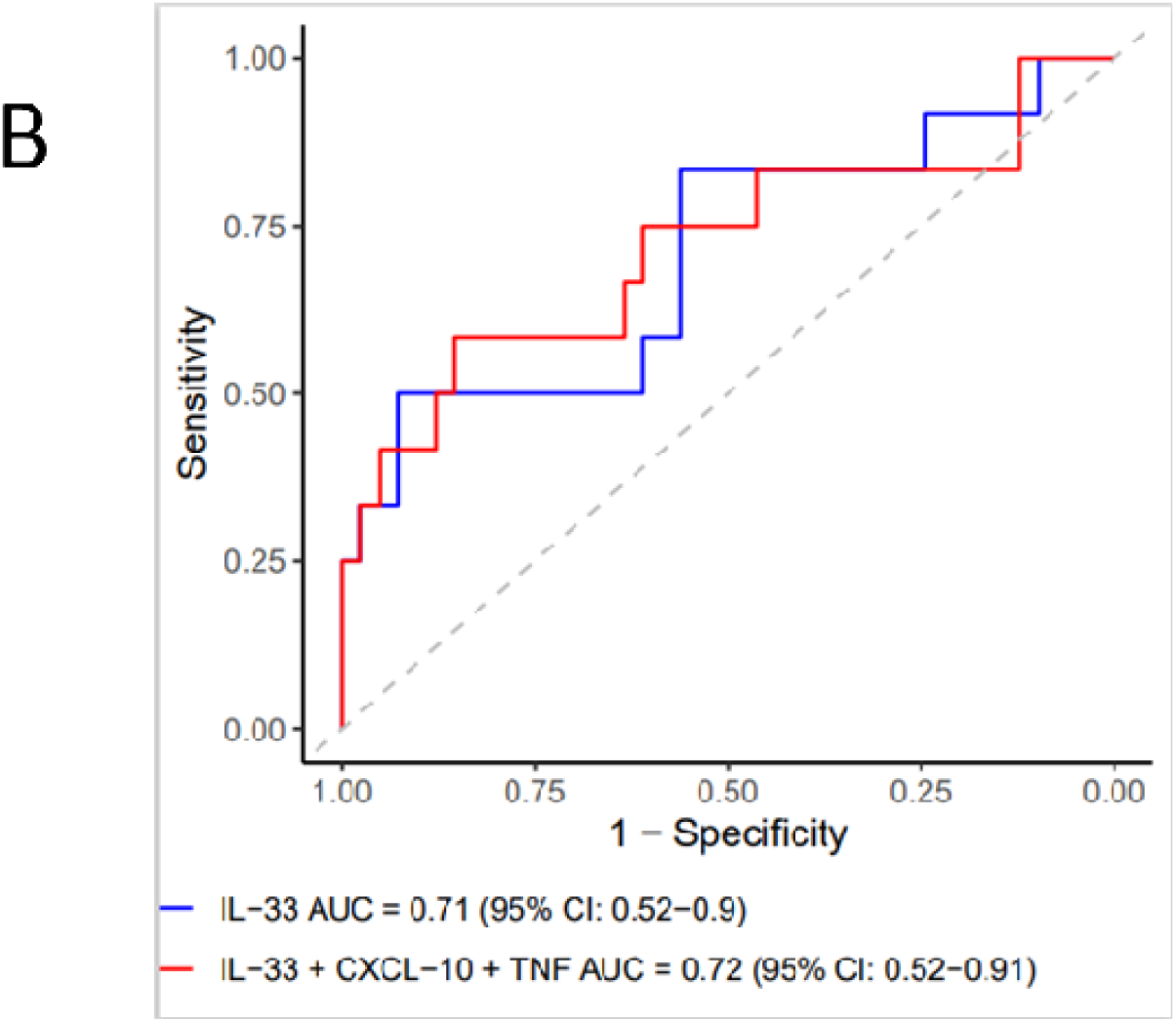
O-link Cytokine Measurements (at index C. difficile infection diagnosis) Compared by Subsequent Uncomplicated versus Recurrent Infection Outcomes. (A)Two-sided Wilcoxon Rank Sum tests were performed comparing cytokines measurements taken within 48 hours of C. difficile diagnosis and compared between patients with an uncomplicated infection (defined as recurrence-free survival by 8 weeks) versus patients who developed a recurrent infection within 8 weeks. (B) ROC Curve Analysis of Univariable and Multivariable Logistic Regression Models for Predicting Recurrent C. difficile infection within 8 weeks. The univariate model shown in blue includes only IL-33 as a predictor, while the multivariable model includes all three cytokines that were significantly altered between patients who did versus did not have recurrence (IL-33, CXCL-10, and TNF). Area under the curve (AUC) values were calculated using the DeLong method with 95% confidence intervals. **, *, ns (non-significant) correspond with P values <0.01, <0.05, and ≥0.05, respectively.

